# Time-resolved analysis of Wnt-signaling reveals β-catenin temporal genomic repositioning and cell type-specific plastic or elastic chromatin responses

**DOI:** 10.1101/2022.08.05.502932

**Authors:** Pierfrancesco Pagella, Simon Söderholm, Anna Nordin, Gianluca Zambanini, Amaia Jauregi-Miguel, Claudio Cantù

## Abstract

Wnt signaling orchestrates gene expression via its effector β-catenin. Whether β-catenin targets genomic regions simultaneously or in a temporal fashion, and how this impacts the chromatin dynamics to modulate cell behavior, is currently unknown. Here we find that β-catenin binds different loci at each time-point after stimulation, implying that the definition of Wnt-targets is fundamentally temporal. This process is intrinsically cell-type specific. In fact, Wnt/β-catenin progressively shapes the chromatin of human embryonic stem cells consistent with their mesodermal differentiation: we call this genomic response plastic. In embryonic kidney cells, on the other hand, Wnt/β-catenin drives a transient chromatin opening, followed by a re-establishment of the pre-stimulation state: a response that we define elastic. Finally, the Wnt-induced transient chromatin opening requires β-catenin, suggesting a previously unappreciated pioneering role for this molecule. We submit that the plastic-vs-elastic behavior constitutes part of the mechanism explaining how Wnt/β-catenin drives divergent cell-fate decisions during development and homeostasis.

## Introduction

Wnt/β-catenin signaling is a highly organized biochemical cascade that triggers a gene expression program in the signal-receiving cell. The Wnt/β-catenin-driven transcriptional response is involved in virtually all cellular processes during development and homeostasis while its deregulation causes human disease (Rim et al., 2022). However, how this response is integrated into lineage-specific choices is still unknown. In the current canonical model of Wnt signaling, the cytoplasmic pool of β-catenin is rapidly depleted by a so-called “destruction complex” that consists of AXIN, the adenomatous polyposis coli (APC) tumor suppressor protein, glycogen synthase kinase 3 (GSK3α and GSK3β, both referred to as GSK3) and casein kinase 1 (CK1) (van Kappel and Maurice, 2017; Wiese et al., 2018). WNT ligands inhibit the destruction complex, causing the accumulation and the nuclear import of β-catenin (MacDonald and He, 2012). In the nucleus, β-catenin associates with the TCF/LEF family of transcription factors, and activates the transcription of Wnt target genes (Mosimann et al., 2009). These mechanisms have been historically studied in several model organisms (Irion and Nüsslein-Volhard, 2022) and cell lines, such as the human embryonic kidney cells (HEK293T) (Li et al., 2012).

Wnt target genes have been collectively defined based on i) their sudden transcriptional activation upon Wnt pathway stimulation (Chang et al., 2008a; Van de Wetering et al., 1997), ii) the presence of Wnt Responsive Elements (WRE) within their genomic loci (e.g. Chang et al., 2008; Jho et al., 2002), or/and (iii) the physical association of TCF/LEF-β-catenin to the DNA in proximity of their annotated transcriptional start sites (TSS) (Doumpas et al., 2019, 2021; Moreira et al., 2018; Schuijers et al., 2014). While these approaches provided us with substantial knowledge about groups of Wnt target genes in different contexts, two outstanding questions remain unanswered. The first concerns time: it is not known whether β-catenin associates with its targets simultaneously or in a time-dependent fashion. For instance, while TCF/LEF and other components of the Wnt transcriptional complex are constitutively associated with the chromatin, it is β-catenin’s arrival, upon Wnt induction, that launches target genes transcription (de la Roche and Bienz, 2007; van Tienen et al., 2017). Therefore, discovering the dynamics of the genome-wide β-catenin binding pattern is required to unambiguously define the direct targets of Wnt signaling. The second question regards how the Wnt/β-catenin cascade modulates chromatin behavior: to date, there exists no comprehensive genome-wide annotation of changing chromatin patterns upon pathway activation. This is important, as shifts in chromatin patterns might underlie how different cells promote diverging gene expression programs in response to Wnt.

To address the first question, here we leveraged on a novel CUT&RUN-LoV-U (Cleavage Under Targets & Release Using Nuclease - Low Volume - Urea) (Zambanini et al., 2022) protocol to establish the full sets of β-catenin target genes detectable over time in two paradigmatic human cell types. We consider CUT&RUN and related technologies particularly powerful for this, as they allow for distinguishing high-confidence direct targets from indirect ones – that is, several genes could be transcriptionally upregulated as secondary or tertiary effects, and WREs could be found *in silico* but might not have functional relevance *in vivo*. This approach allowed us to establish that β-catenin repositions to different genomic loci along stimulation time, showing that a definition of Wnt target genes must take into account the time-dimension. Moreover, β-catenin physical targets are largely cell-type specific, as only a subset of them are present across the examined contexts. Of note, we were surprised in observing that β-catenin binding is associated with both activation and repression of cell-specific gene expression programs, underscoring how Wnt/β-catenin drives complex cell behaviors.

Second, to mechanistically understand the causal relation between β-catenin binding and the induced genetic programs, we analyzed how this pathway orchestrates the chromatin dynamics genome-wide across defined time points. We found that human embryonic stem cells (hESCs) respond to Wnt/β-catenin activation by progressively shaping their chromatin accessibility profile in a manner that is consistent with their gradual acquisition of a mesodermal identity: differentiation genes loci open over time, while pluripotency ones close. We refer to this genomic response as *plastic*. On the other hand, human embryonic kidney cells (HEK293T), which are known to be highly responsive to Wnt activation (Gujral and Macbeath, 2010) (Li et al., 2012), appear more resistant to a long-term Wnt/β-catenin-driven change in cell identity. In this context, the chromatin displays a temporary opening of relevant regions at 4 hours after stimulation, followed by a re-establishment of its pre-stimulation state: we define this transient response as *elastic*. Finally, by using genetic tools and enzymatic inhibitors, we demonstrate that the transient chromatin opening mechanistically requires β-catenin together with histone acetyltransferase and deacetylase enzymatic activities, unearthing a previously overlooked chromatin pioneering function of β-catenin. We propose that the plastic and elastic responses represent modalities of genomic behaviors induced by Wnt/β-catenin that underlie how this signaling pathway can elicit radically divergent responses depending on the cellular context.

## Results

### A time-resolved atlas of β-catenin genome-wide physical occupancy

We set out to establish the genome-wide dynamics of β-catenin binding over time, in two paradigmatic human cells, HEK293T and hESCs. Here, Wnt/β-catenin signaling can be reliably activated by administration of the GSK3 inhibitor CHIR99021 (hereafter CHIR), which induces β-catenin stabilization, nuclear translocation, and activation of downstream targets (Figure 1A) (Doumpas et al., 2021; Wray et al., 2011). We applied CHIR for up to 3 consecutive days and performed CUT&RUN-LoV-U (Zambanini et al., 2022) targeting β-catenin at selected time-points after Wnt pathway stimulation (Figure 1B). This approach generated a time-resolved, cell-specific compendium of the genome-wide β-catenin chromatin occupancy profiles (Table S1).

**Figure 1.**
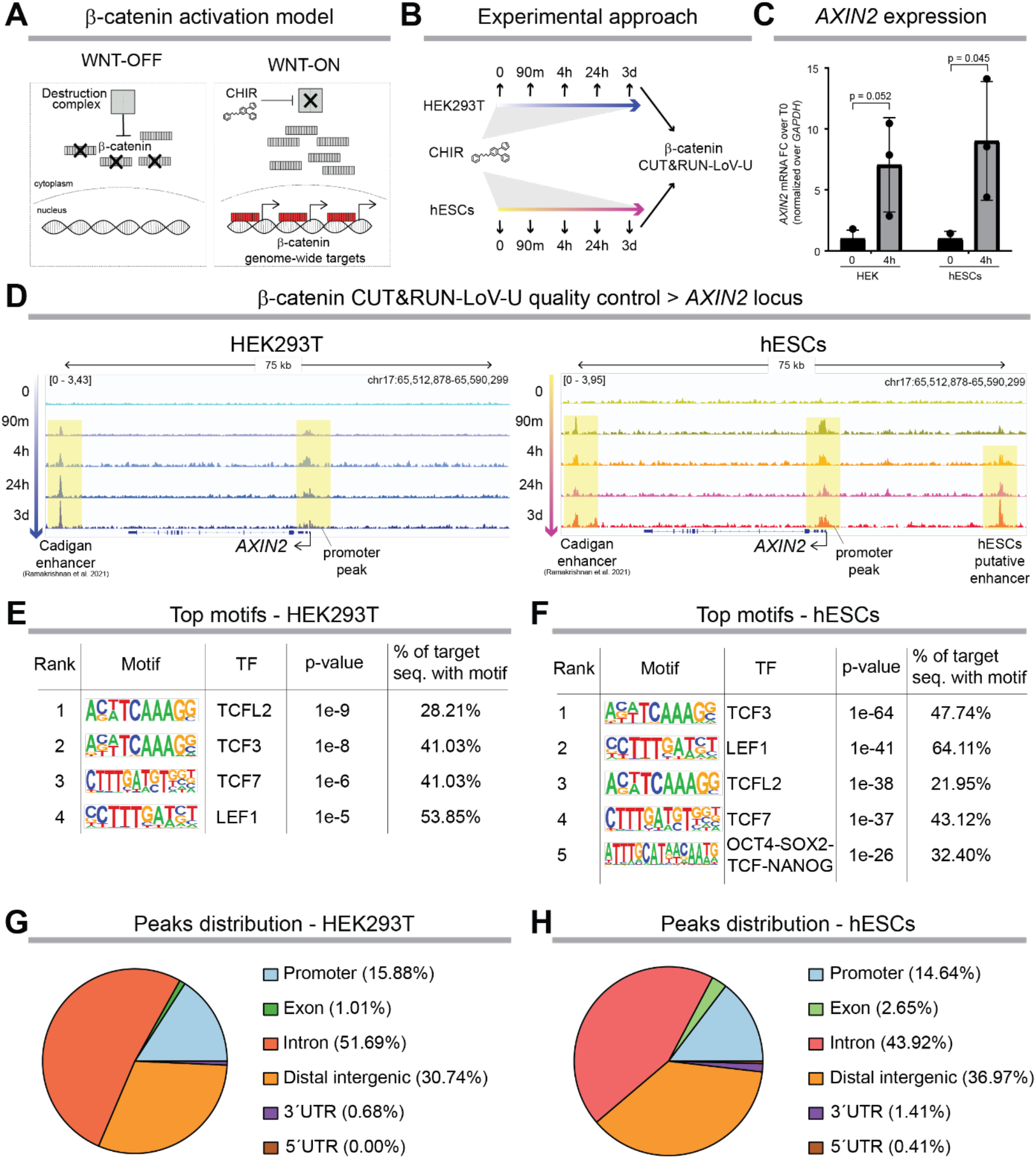
A time-resolved atlas of β-catenin genomic occupancy across two human cell types. (A) Wnt/β-catenin signaling was activated by treating cells with the GSK3 inhibitor CHIR99021 (CHIR). CHIR inhibits the destruction complex, leading to the accumulation of cytoplasmic β-catenin, which in turn translocates to the nucleus. Here, β-catenin reaches specific genomic locations where it interacts with TCF/LEF transcription factors and activates the expression of Wnt target genes. (B) Experimental approach: HEK293T and hESCs were treated with CHIR for up to 3 days. Cells were collected at 90 minutes, 4 hours, 24 hours, and 3 days after Wnt activation, and processed for CUT&RUN-LoV-U against β-catenin. (C) Upregulation of the mRNA expression of paradigmatic Wnt target genes, e.g., AXIN2, confirmed the activation of the Wnt/β-catenin signaling pathway already after 4 hours of CHIR treatment. (D) Reproducible (n = 3 independent experiments; IDR as described in methods section) β-catenin peaks were observed at Wnt target genes loci. We could consistently observe β-catenin peaks in correspondence of the AXIN2 promoter, starting already at 90 minutes after CHIR treatment and proceeding for 3 days. We confirmed the binding of β-catenin on a downstream enhancer, that we term “Cadigan enhancer”, identified by (Ramakrishnan et al., 2021). We further report a hESCs-specific β-catenin binding event approximately 30 kb upstream of the AXIN2 TSS, which could indicate a putative hESCs-specific regulatory region. (E) Motif analysis (HOMER) performed on β-catenin peaks called in HEK293T showed preponderance of TCF/LEF binding sequences. (F) Motif analysis of β-catenin peaks in hESCs confirmed TCF/LEF as the most abundant transcription factor, followed by pluripotency players – SOX2, POU5F1/OCT4, NANOG - –, suggesting a potential functional interaction between β-catenin/TCF/LEF and these key transcriptional regulators, in accordance with previous work in embryonic stem cells (Kelly et al., 2011). (G, H) Both in HEK293T and hESCs, β-catenin peaks were primarily localized within intronic and distal intergenic genomic regions. A similar fraction of β-catenin binding events occurred in correspondence of promoters (14-15%), while the fraction of peaks localized on exon regions was double in hESCs (2.65%) than in HEK293T (1.01%). Abbreviations: FC: fold change; HEK = HEK293T, i.e., human embryonic kidney cells 293T; hESCs: human embryonic stem cells; TF: transcription factor.(Kelly et al., 2011)

Key analyses support the reliability of our approach. First, CHIR treatment caused a comparable upregulation of the Wnt pathway in both cell types, monitored by *AXIN2* mRNA expression (Figure 1C). Second, CUT&RUN-LoV-U generated reproducible β-catenin peaks in previously characterized WREs in correspondence to the *AXIN2* genomic locus at all time points (Figure 1D) and in several other loci, comparable to previous datasets (Figure S1). Third, both cell types display statistical enrichment for motifs of all four individual TCF/LEF proteins as primary transcription factor footprinting in the sequences underlying β-catenin peaks (Figure 1E, F). Finally, these datasets recapitulate the previously observed genomic distribution of β-catenin in Wnt-ON conditions across functionally diverse regions and with respect to average distance to annotated genes (Figure 1G, H) (Cantù et al., 2018; Zambanini et al., 2022). Notably, our time-resolved CUT&RUN-LoV-U dataset robustly reproduced the β-catenin binding patterns observed via single-time-point ChIP-seq (performed after 24 hours of CHIR stimulation) in a previous report (Doumpas et al., 2019). However, single-time-point ChIP-seq failed to detect time-restricted β-catenin binding events that occur transiently at other individual time-points (Figure S1). Overall, these datasets represent a comprehensive comparison of β-catenin physical genome-wide occupancy, over time, and in different cellular models.

### β-catenin physical targets change over time and are largely cell-type specific

We used highly stringent peak calling parameters and considered only binding events that occur reproducibly in our biological replicates to converge on sets of high confidence β-catenin physical targets at each time point. Notably, β-catenin targets differ at each time point in HEK293T and hESCs both in number and identity (Figure 2A, B). We generated a timecourse recording of β-catenin “movements” on the genome to isolate those instances that correspond to time-point-specific regulation events (Figure 2B). Several genes are targeted by β-catenin as early as 90 minutes after pathway stimulation, while others require unexpectedly longer stimulation times, including some that were only detected at day 3 (Figure 2C, right panels). Among these, in HEK293T we scored the relevant *E2F7*, recently identified as Wnt/TBX3 target repressed to regulate mitosis (Jin et al., 2022). Our analysis suggests that the regulation of the cell cycle by Wnt signaling might occur, in cascade, among the late events. In hESCs, the early-time specific regulation of pluripotency factors, such as *NANOG*, is followed by targeting other genes, including *VGLL4*, which acts as a positive promotor of hESCs survival (Tajonar et al., 2013) and negative feedback regulator of Wnt signaling (Jiao et al., 2017).

**Figure 2.**
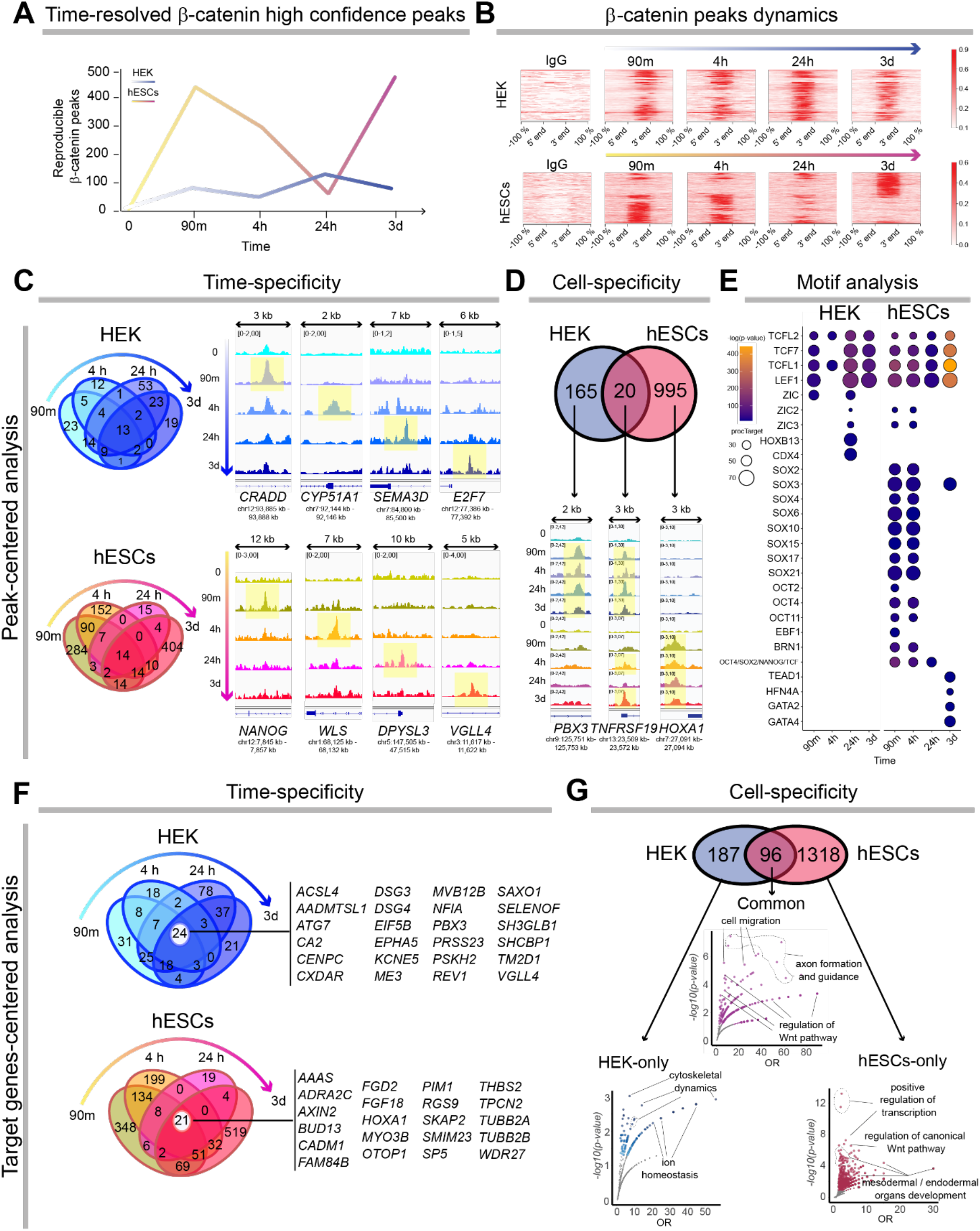
β-catenin physical targets are time-dependent and context-specific. (A) Reproducible β-catenin binding events were detected in different numbers across the selected time points and cellular contexts. (B) The enrichment heatmaps represent the reproducible β-catenin peak dynamics at each time point (from left to right) in the two cell types. The peaks signal intensity is centered at the peak summit for the corresponding locus. Each horizontal line across the charts represents one genomic location, emphasizing that the same position might bear a signal at one but not at other time-points. (C) The majority of β-catenin peaks are time-dependent. Only 13 peaks in HEK293T and 14 peaks in hESCs are consistently bound by β-catenin during Wnt activation. In HEK293T, late-only targets include cell cycle regulators such as E2F7. In hESCs, β-catenin initially targets several loci involved in the control of pluripotency, such as NANOG. At successive time points, β-catenin binds to regions associated with different classes of genes, including modulators of the Wnt signaling pathway itself (e.g., WLS, VGLL4). (D) β-catenin peaks are mostly cell-specific. We identified only 20 peaks shared between HEK293T and hESCs. Common targets include well-characterized Wnt target genes, such as AXIN2 and TNFSRF19, while those that are cell-specific reflect key processes driven by Wnt in the two cell types. (E) Dot blot shows HOMER-based motif analysis performed using the sequences underlying β-catenin peaks. This identified TCF/LEF consensus sequences as those most frequently represented, both in HEK293T and hESCs. HEK293T also showed enrichment of CDX consensus sequences. In hESCs, β-catenin peaks were associated with motifs for pluripotency factors (e.g., SOX and POU/OCT families of transcription factors) during the early phases of Wnt activation; at later time points, peak-associated-regions were enriched with motifs for transcription factors involved in mesodermal differentiation (e.g., GATA). (F) Peaks were assigned to genes using GREAT (McLean et al., 2010). Only a small number of genes are constitutively targeted by β-catenin over time (24 in HEK293T, 21 in hESCs). (G) HEK293T and hESC share less than 100 β-catenin target genes, and these are involved in the modulation of basic processes such as cell migration as well as the regulation of the Wnt pathway itself. hESCs-specific β-catenin targets are broadly involved in the differentiation towards the meso-endodermal lineages, while the fewer HEK293T-specific targets are mostly associated with cytoskeletal dynamics and the modulation of ion homeostasis. All genomic locations are approximated to the kilobase (kb) (McLean et al., 2010).

Overall, only a fraction of the β-catenin binding events occurs consistently over time, while the major peak subsets are consistently time-point specific (Figure 2C); this reveals the transient nature of β-catenin association to the chromatin. The comparison between HEK293T and hESCs uncovered that the vast majority of all β-catenin binding events, notwithstanding the time at which they occur, are cell-type specific (Figure 2D). While previously identified targets, such as *AXIN2* (Jho et al., 2002; Figure 1) and *TNFRSF19/TROY* (Fafilek et al., 2013; Figure 2D), were present in both groups, considerably wider were the sets of HEK293- and hESCs-specific target genes (Figure 2D, Figure S2).

Our experiment therefore exposes genes that are regulated by Wnt/β-catenin signaling in a cell type-specific fashion and provides a reference dataset to investigate the mechanisms of this regulation. As examples, we performed two analyses that exemplify the versatility of our dataset. First, orthogonal motif analysis that considers both time- and cell-resolved data points (Figure 2E) permitted us to draw the following conclusions: i) the predominant, if not unique, response of Wnt/β-catenin to GSK3 inhibition in HEK293T is mediated by TCF/LEF transcription factors, in agreement with previous studies (Schuijers et al., 2014); ii) contributions of other transcription factors, such as the CDX homeobox proteins (Ramakrishnan et al., 2021), could fine-tune this response; iii) in hESCs, the TCF/LEF footprint is accompanied by motifs for OCT4 (Kelly et al., 2011) and those of other pluripotency factors (Cole et al., 2008), followed by a gradual appearance of GATA consensus sequences that mark mesoderm differentiation (Molkentin et al., 1997). As a second analysis, we searched for the biological processes underlying β-catenin time-course cell-specific behavior by generating a peak-to-gene(s) annotation based both on proximity and functional characterization of regulatory regions using GREAT (McLean et al., 2010), to then assign cell-specific biological processes that emerge across time after stimulation (Figure 2F, G, Figure S2). Overall, Wnt target genes in HEK293T are involved in cell proliferation, motility, and neural events, consistent with the ambivalent renal/neural nature of these cells (Figure 2G, Figure S2) (Lin et al., 2014; Shaw et al., 2002). hESCs, on the other hand, upon Wnt/β-catenin activation, progressively regulate genes that are associated with the exit from pluripotency and commitment towards mesodermal lineage and heart development (Figure 2G, Figure S2).

Collectively, these analyses emphasize that the CHIR-mediated Wnt/β-catenin stimulation is sufficient to drive different outcomes depending on the nature of the signal-receiving cell.

### Differential transcription changes are associated with the stage-specific binding patterns of β-catenin

The time- and cell-type-specific genomic association of β-catenin observed suggests that Wnt/β-catenin signaling regulates transcription over time. To test this, we aimed at associating each β-catenin binding event with mRNA expression changes of the peak-annotated-genes. We generated datasets to include the global transcriptional response of Wnt-activated HEK293T and hESCs by using bulk RNA sequencing (Figure S3). By merging the differential expressed genes (DEGs) with the peak-associated gene lists present at any time point, we were able to identify HEK293T-specific, hESCs-specific, and shared DEGs. The shared group included the positive regulation of well-characterized key Wnt target genes such as the previously mentioned *AXIN2, DKK1, NKD1* and, *TNFRSF19/TROY* (Figure 3A). The HEK293T and hESC groups of upregulated DEGs, on the other hand, comprised factors that are known to have tissue-specific regulation, such as the developmental transcription factor *TBX3* in HEK293T and the mesoderm/cardiac-relevant *MSX1* and *NODAL* in hESCs (Figure 3A, left and right heatmaps). In all the three groups, we identified a considerable fraction of β-catenin-bound (hence, direct targets) genes that are transcriptionally downregulated (Figure 3A, blue bars and signal): this exposes, in a quantitative fashion, a previously unappreciated negative effect on transcription promoted by Wnt/β-catenin in both cell types. Notable among the hESCs-specific β-catenin-bound downregulated DEGs, we scored *SOX2, NANOG* and other genes encoding for stemness or pluripotency factors (Ying et al., 2008). Whilst others have shown that Wnt/β-catenin contributes to the exit of hESCs from pluripotency (Davidson et al., 2012), our results further suggest that it might do so by acting as a direct transcriptional repressor of the pluripotency circuit.

**Figure 3.**
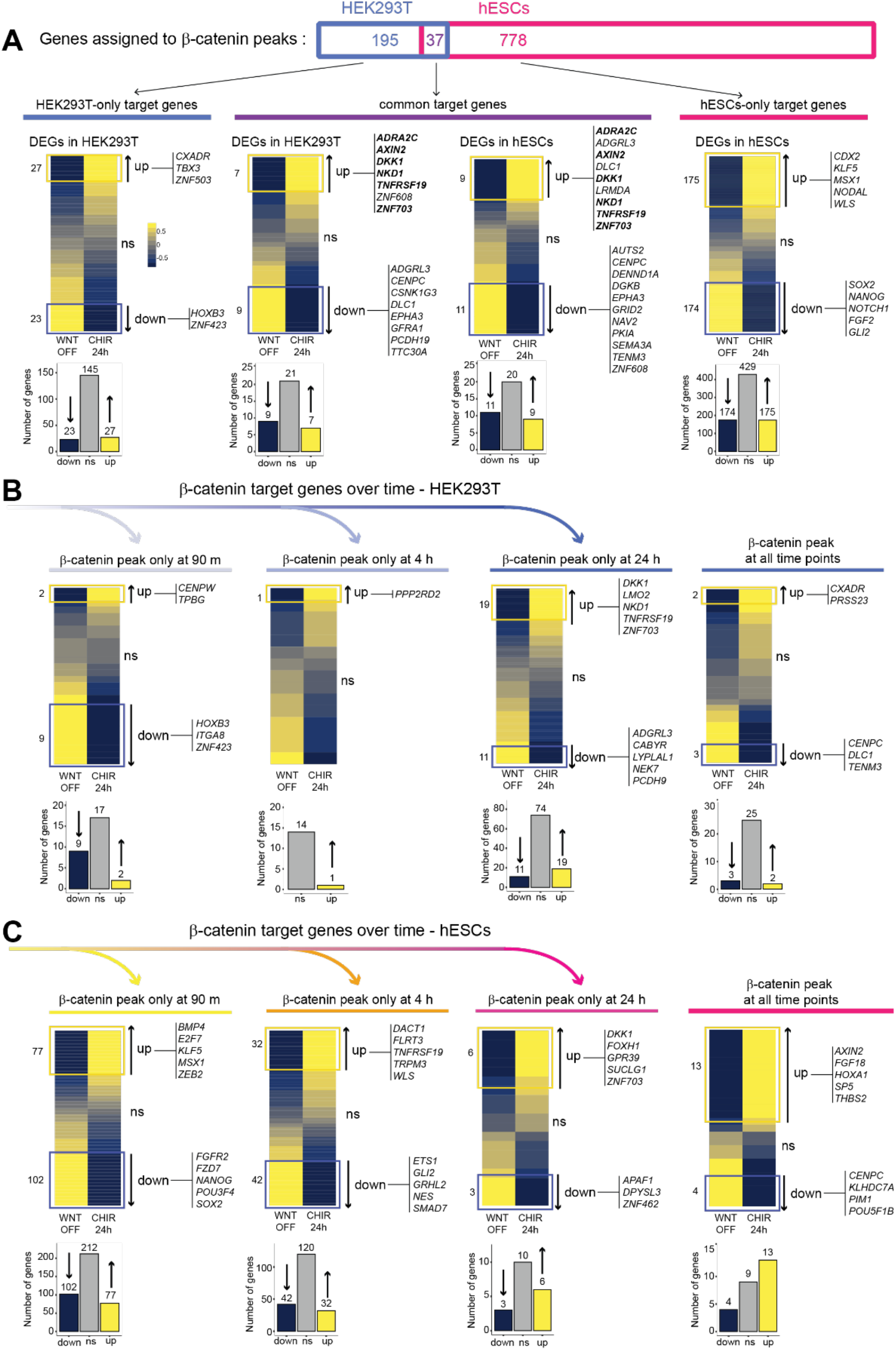
Differential β-catenin binding is associated with both upregulation and downregulation of transcription. (A) We selected genes physically bound by β-catenin within the first 24 hours of Wnt activation (90 minutes, 4 hours, 24 hours CUT&RUN-LoV-U time-points) and assessed their transcriptional response by RNA sequencing (n = 3 for each condition and cell type). Heatmaps provide overviews of significantly upregulated genes (yellow box, upwards arrow, “up” label), significantly downregulated genes (blue box, downwards arrow, “down” label), and non-affected genes (“ns” label) among HEK-specific, HEK-hESC-common, and hESCs-specific β-catenin targets (from left to right, respectively). The corresponding histograms summarize the exact numbers of upregulated, downregulated, and non-affected β-catenin targets. Among the common targets, the well-characterized Wnt target genes *AXIN2, NKD1, DKK1, TNFRSF19* and *ZNF703* were all significantly upregulated upon β-catenin binding both in HEK293T and hESCs (gene names marked in bold font). Notably, in hESCs β-catenin interaction with loci coding for key pluripotency factors (*SOX2, NANOG*) was associated with their downregulation. (B) Time-resolved analysis of the correlation between β-catenin peaks and gene expression changes in HEK293T. Most β-catenin binding events do not correspond to alterations in the expression of the gene assigned to the peak. Genes targeted at 90 minutes show a stronger tendency to be downregulated (left heatmap), while later targets display a higher tendency to be upregulated. (C) Time-resolved analysis of the correlation between β-catenin peaks and gene expression changes in hESCs. Early targets (90 minutes) include pivotal genes involved in the exit from pluripotency. For example, *SOX2, NANOG* and *FGFR2* are all targeted transiently by β-catenin and their expression significantly downregulated. A comparable transient association on the loci encoding *KLF5, MSX1, ZEB2*, and *BMP4* results in their significant upregulation (left heatmap). An analogous balance between downregulation and upregulation of β-catenin targets is also observed at intermediate targets (4 hours), while loci targeted consistently by β-catenin tend to be upregulated (right heatmap).

We then crossed the transcriptional changes with the time-resolved dynamics of β-catenin genomic binding, to resolve how the abovementioned genes and processes are temporally regulated (Figure 3B, C). While HEK293T displayed a low number of time-dependent changes in both the up- and down-regulation directions consistent with a refractory tendency in changing identity (Figure 3B), hESCs appeared more affected and presented, especially in the initial phases of stimulation, downregulation of pluripotency genes, such as *SOX2* and *NANOG*, accompanied by upregulation of transcription factors like *KLF5*, that likely contributes to retard mesodermal differentiation in the early phases after stimulation (Aksoy et al., 2014). At later time-points, hESCs appear to consolidate the mesoderm>cardiac fate by activating, for example, the gastrulation-relevant, mesoderm-patterning *SP5* (Weidinger et al., 2005) and the heart morphogenesis factor *HOXA1* (Roux et al., 2015).

This time-course analysis converges with the conclusions drawn by our timed peak-gene annotations (Figure 2C-E) and the time-lapse appearance of regulated biological processes (Figure 2G, S2) to uncover the cell-type specific gene regulation operated by Wnt/β-catenin signaling, and how this affects cell behavior of different cell types.

### Wnt/β-catenin remodels the chromatin accessibility landscape over time in a cell-specific manner

Chromatin accessibility emerged as a golden standard to assess the causal relation between binding of transcriptional regulators (Figure 2) and the implementation of transcriptional outputs (Figure 3) (Agbleke et al., 2020). To date, the time-dependent changes in chromatin landscape imposed by Wnt/β-catenin induction have not been resolved (Figure 4A). To this end, we harvested HEK293T and hESCs at the relevant time points after pathway induction and performed Assay for Transposase-Accessible Chromatin using sequencing (ATAC-seq; Buenrostro et al., 2013) in triplicate per time-point (Figure 4B, Figure S4). This yielded reproducibly changing patterns of chromatin accessibility profiles that landmark the different time points: these include signals in the proximity of Wnt targets (e.g., *AXIN2, SP5*), pluripotency (e.g. *OCT4*), and differentiation-relevant factors (e.g., *TBXT/BRA*) in triplicate per time-point (Figure 4B, Figure S4, Table S2). HEK293T and hESCs behaved in a radically different manner upon Wnt/β-catenin activation. While HEK293T displayed relatively lower fluctuations in the number of regulated chromatin regions (Figure 4C, green>blue components in the charts), consistent with the low number of β-catenin peak-associated transcriptional changes (Figure 3B), hESCs showed a consistently higher number of open loci in their ground state, suddenly followed (4 h) by a massive chromatin closure (corresponding to a ca. 40% reduction; Figure 4C, yellow>red components in the charts). At later time-points we recorded opening of regulatory regions associated with the action of lineage-specific transcription factors: among these, the depletion of regions containing POU and NANOG motifs, and enrichment of those for the AP1 complex, TEAD, FOS, and SMAD transcription factors (Figure 4D). This recapitulates the well-known phenomenon by which the global transcriptional hyperactivity of hESCs is reflected in a wide opening of the chromatin (Efroni et al., 2008), while their differentiation requires genomic specialization via wide-spread closure and positive regulation of select regions by lineage-specific factors (Orkin and Hochedlinger, 2011). Our observation implies that sustained Wnt/β-catenin activation is sufficient to drive this process (Figure 4C).

**Figure 4.**
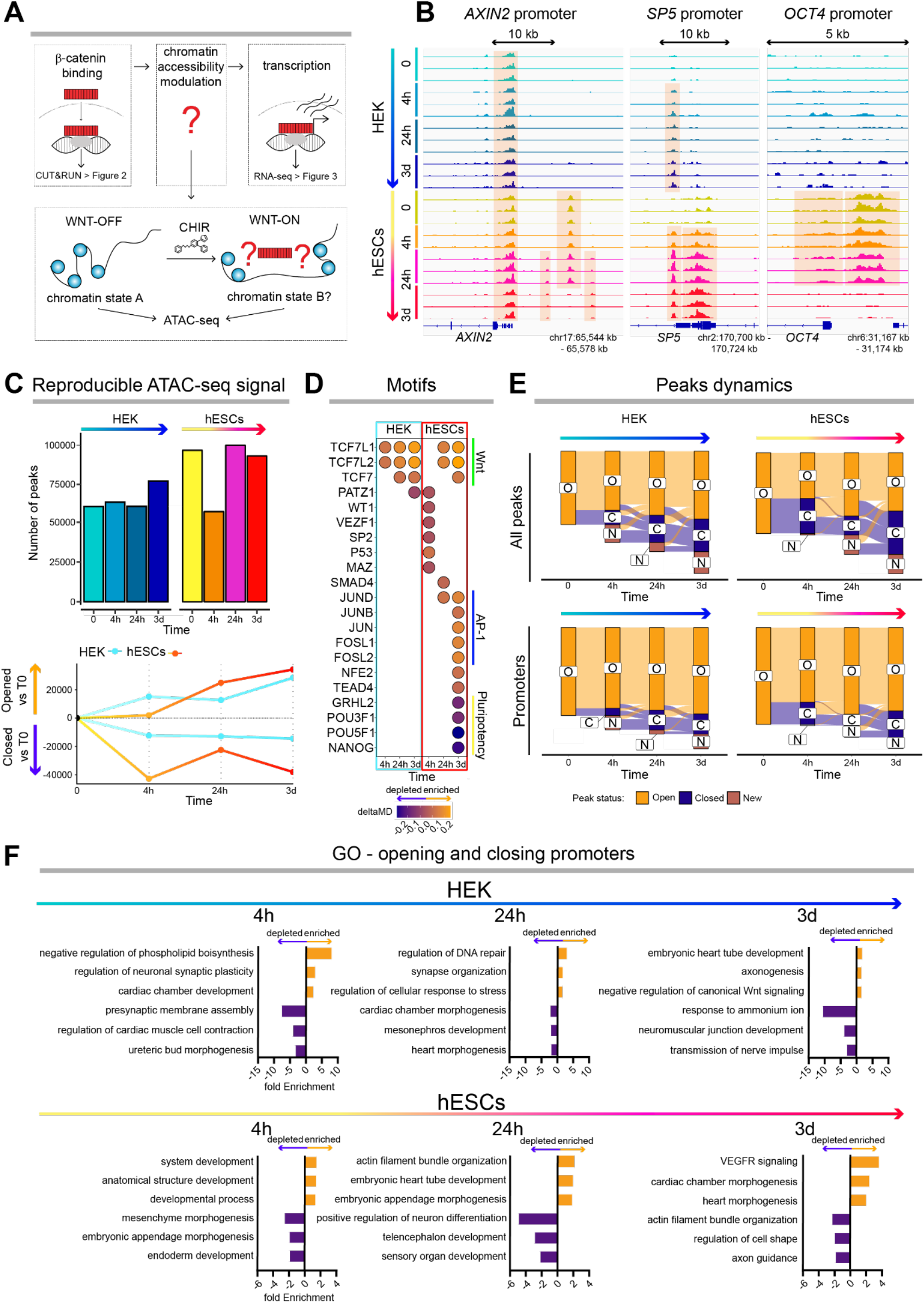
Wnt/β-catenin signaling modulates genome-wide chromatin accessibility in a time-resolved and cell-specific manner. (A) Upon Wnt signaling activation, β-catenin binds specific target loci and modulates their transcription. Regulation of gene expression requires changes in chromatin accessibility, yet whether and how Wnt/β-catenin affects chromatin dynamics has not been thoroughly investigated. (B) Temporal dynamics of Wnt/β-catenin chromatin response by performing ATAC-sequencing profiles in HEK293T and hESCs treated with CHIR for 4 hours, 24 hours and 3 days (n = 3 per condition, for each cell type). Shown are the chromatin accessibility profiles that trace marked temporally-resolved dynamics in correspondence of Wnt target genes (AXIN2, SP5) and genes coding for pluripotency factors (POU5F1/OCT4). (C) Temporal dynamics of genomewide chromatin accessibility (ATAC-sequencing peaks). HEK293T cells show a moderate and homogeneous response to Wnt/β-catenin activation; at every time point, we could observe a similar number of newly opened and newly closed regions (approx. 10’000 peaks). hESCs display a more radical and directional response: already at 4 hours, more than 40’000 peaks disappeared, while a very limited number of regions opened, a pattern consistent with the known global chromatin closure that accompanies the exit from pluripotency (Orkin and Hochedlinger, 2011). Conversely, over 20’000 new regions open at 24 hours, and further high numbers of new regions open or close after 3 days. (D) Motif analysis of peaks appearing (motifs enriched; orange side of the color spectrum) and disappearing (motifs depleted; violet side of the color spectrum) at each time point. Wnt/β-catenin induces a TCF/LEF-restricted response in HEK293T, as at every time point TCF/LEF motifs are the only ones to be enriched within newly accessible regions. In hESCs, the first response is accompanied by enrichment for P53 and SP motifs, and only after 24 hours are TCF/LEF motifs enriched. After 3 days of stimulation, motifs for pluripotency factors (e.g., POU5F1 and NANOG) are depleted, while motifs for non-Wnt pathway effectors are enriched (e.g., JUN, FOS, TEAD). (E) Alluvial plots show the ATAC-sequencing-peaks temporal dynamics. Many regions undergo consecutive rounds of chromatin opening and closing over time. (F) Gene ontology (GO) – biological processes analysis (GO) of genes assigned to promoter regions that became accessible or inaccessible at different time points from Wnt/β-catenin activation (selected GO terms, p < 0.05, fdr < 0.05). In HEK293T, Wnt/β-catenin induces a rapid closure of regions coding for genes associated with renal identities, while dynamically modulating loci involved in neural differentiation – in accordance with the ambiguous kidney/neural identity of HEK293T cells (Lin et al., 2014). In hESCs, many loci involved in both pluripotency maintenance and cell differentiation were rendered inaccessible in the first 4 hours. At later time points, promoters associated with mesodermal differentiation, and specifically with heart development, were progressively opened. All genomic locations are approximated to the kilobase (kb).

To pinpoint which are the relevant genes associated with the chromatin changes, we used ChIPseeker (Yu et al., 2015) to filter all the hits falling within annotated promoters. We noticed that, when considering ATAC peak signals that appear or disappear, promoters show a lower dynamicity in their behavior (Figure 4E, lower panels) compared to the global genomic responses (Figure 4E, upper panels). This is consistent with the idea that individual promoters can be regulated by multiple tightly-regulated developmental enhancers (Zabidi et al., 2015). Promoter-associated genes were grouped based on their opening or closing behavior at each time point in the two cell types (Figure 4F). HEK293T, consistent with their ambivalent nature at the interface between embryonic kidney and neural tissues (Lin et al., 2014; Shaw et al., 2002), regulate promoters of genes relating to synaptic plasticity, ureteric bud morphogenesis and mesonephros development (Figure 4F, upper bar-plots). Notably, Wnt/β-catenin drives a loss of mesonephros- and ureteric bud morphogenesis-associated genes, in favor of a gain in synapse and neuronal plasticity phenotypes. This is a key observation, as it shows that HEK293T display one of the possible physiologically informative responses to Wnt/β-catenin activation (Baker et al., 1999; Moon and Kimelman, 1998). hESCs, more predictably and in concordance with our previous analyses, respond by closing, within the first 4 hours, the promoters of genes associated with general developmental processes, such as mesenchyme and appendage morphogenesis, to activate, at the successive time-points, genetic programs that are pointing to the attainment of a mesodermal fate and subsequent differentiation into the cardiac lineage (Figure 4F, lower bar-plots). Of note, this is achieved by Wnt/β-catenin without the concomitant addition of the several ingredients typically added to the hESCs cardiogenic medium (Den Hartogh et al., 2015).

Taken together, these analyses revealed how distinct chromatin landscape identities can be patterned by the same signaling pathway to produce diverging cell behaviors.

### Wnt/β-catenin elicits plastic or elastic genomic responses at Wnt target loci

When looking at the changing chromatin profile on the promoter of the paradigmatic target gene *AXIN2*, we noticed that its behavior was different between the two cell types – despite *AXIN2* being a common target (Figure 4B). While Wnt/β-catenin activation induces a quantitative, gradual opening of this region in hESCs, HEK293T react by opening *AXIN2* promoter at 4 hours, to then restore the initial chromatin conformation at 24 hours. Quantification of the differential accessibility across three replicates showed that this behavior was shared by the promoters of other common, notable Wnt target genes such as *DKK1, ISL1* and *LEF1* (Figure 5A). We then looked at the chromatin profile over time of a broader, curated list of genes that include several historically-characterized Wnt targets (https://web.stanford.edu/group/nusselab/cgi-bin/wnt/target_genes; Figure 5B, Figure S5). All these showed the same dichotomic pattern of response between the two cell types and displayed a strikingly comparable accessibility variation range (Figure 5B).

**Figure 5.**
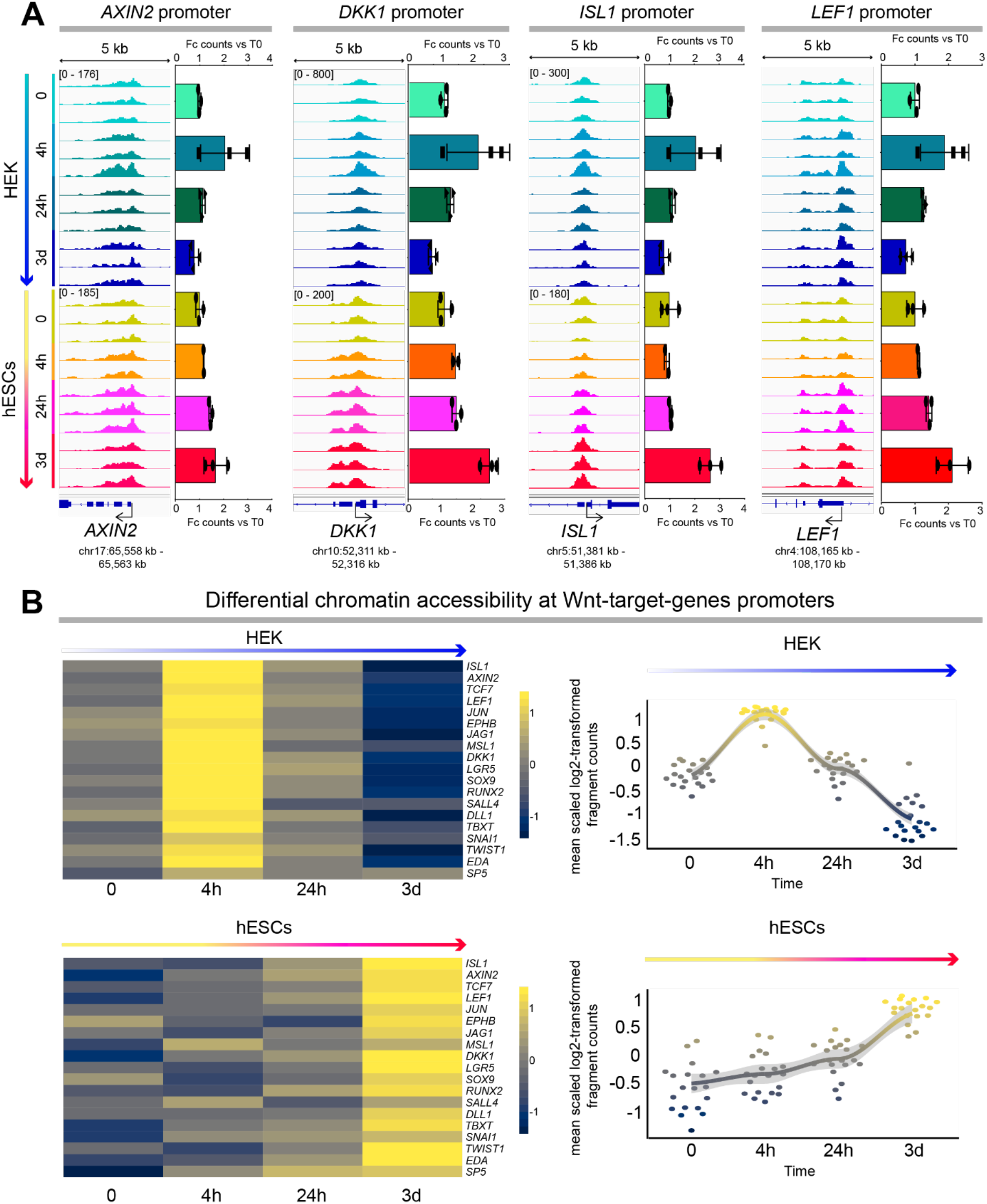
Wnt/ β-catenin signaling elicits plastic and elastic chromatin responses at target gene promoters. (A) Quantification of time-resolved differential chromatin accessibility in correspondence of Wnt target genes promoters in HEK293T (above in green>blue) and hESCs (below in yellow>red) treated with CHIR. In HEK293T, promoters of key Wnt target genes such as *AXIN2, DKK1, ISL1* and *LEF1* show increased chromatin accessibility after 4 hours of CHIR stimulation; within 24 hours, accessibility returns to pre-stimulation levels. We call this response elastic. In hESCs, the same promoters progressively open during the entire duration of Wnt/β-catenin stimulation. We call this response plastic. The values displayed represent fragment-counts (at the promoters-associated peaks) fold-change against time 0. (B) Quantification of differential chromatin accessibility analysis extended to a broader selection of Wnt target genes. Abbreviations: Fc: fold change. All genomic locations are approximated to the kilobase (kb).

We reasoned that the behavior of hESCs is not surprising, as Wnt targets include several genes that are gradually upregulated to shape hESCs’ differentiation fate, such as the Wnt-inducible epithelial-to-mesenchymal-transition factors *SNAI1* (Dias et al., 2020) and *TWIST1* (Reinhold et al., 2006), and the mesoderm driver *TBXT*/*BRA* (Martin and Kimelman, 2010) (Figure 5B, lower panels). As an analogy to the ability to change cell identity, a feature termed plasticity typical of ESCs (Marks et al., 2012), we refer to this genomic response as *plastic*. On the contrary, HEK293T display a seemingly opposite response curve (Figure 5B, upper panels). The considerable opening at 4 hours, followed by the closing at 24, is suggestive of a reversible reaction to restore the initial chromatin state: we refer to this genomic response as *elastic*. This elastic genomic response is consistent with the apparent resilience shown in our temporal peaks-versus-transcriptomics analysis by HEK293T (Figure 3B): we speculate that this represents the refractory abilities of HEK293T in changing cell identity.

Overall, time-resolved ATAC-seq upon Wnt/β-catenin pathway activation shows that the chromatin accessibility patterns at Wnt target genes strictly depend on the cellular context. Moreover, our data unearths new kinetics of chromatin accessibility regulation that underlie gradual lineage commitment as opposed to rapid but transient induction of gene expression programs.

### β-catenin pioneers WREs by triggering chromatin remodeling enzymatic complexes

We set out to determine how Wnt signaling activation triggers the rapid opening of the chromatin in correspondence to Wnt targets’ promoters. The current model considers that β-catenin nuclear entry is preceded by the presence, on WREs, of an already assembled transcriptional complex (van Tienen et al., 2017), and that β-catenin would associate with regions that have been previously rendered permissive by chromatin accessibility and protein condensates (Zamudio et al., 2019). However, this does not explain how CHIR can modulate chromatin opening only after β-catenin becomes stabilized (Figure 5). Therefore, we tested the requirement of the β-catenin protein itself for this process. We employed HEK293T cellular clones knock-out (KO) for β-catenin that we previously generated (Doumpas et al., 2019) and aimed at measuring the chromatin accessibility response upon CHIR treatment (Figure 6A, left panel). We repeated the ATAC profiling and used ATAC Primer Tool (Yost et al., 2018) to quantify the relative signals over stimulation time at Wnt target promoters. While CHIR-treated parental HEK293T reproduced the elastic chromatin response on *AXIN2* and *DKK1* promoters also via this orthogonal, independent measurement, β-catenin KO cells consistently failed in opening these promoter regions, demonstrating that β-catenin is necessary to induce the chromatin opening response observed at 4 hours after CHIR stimulation (Figure 6A, right panels).

**Figure 6.**
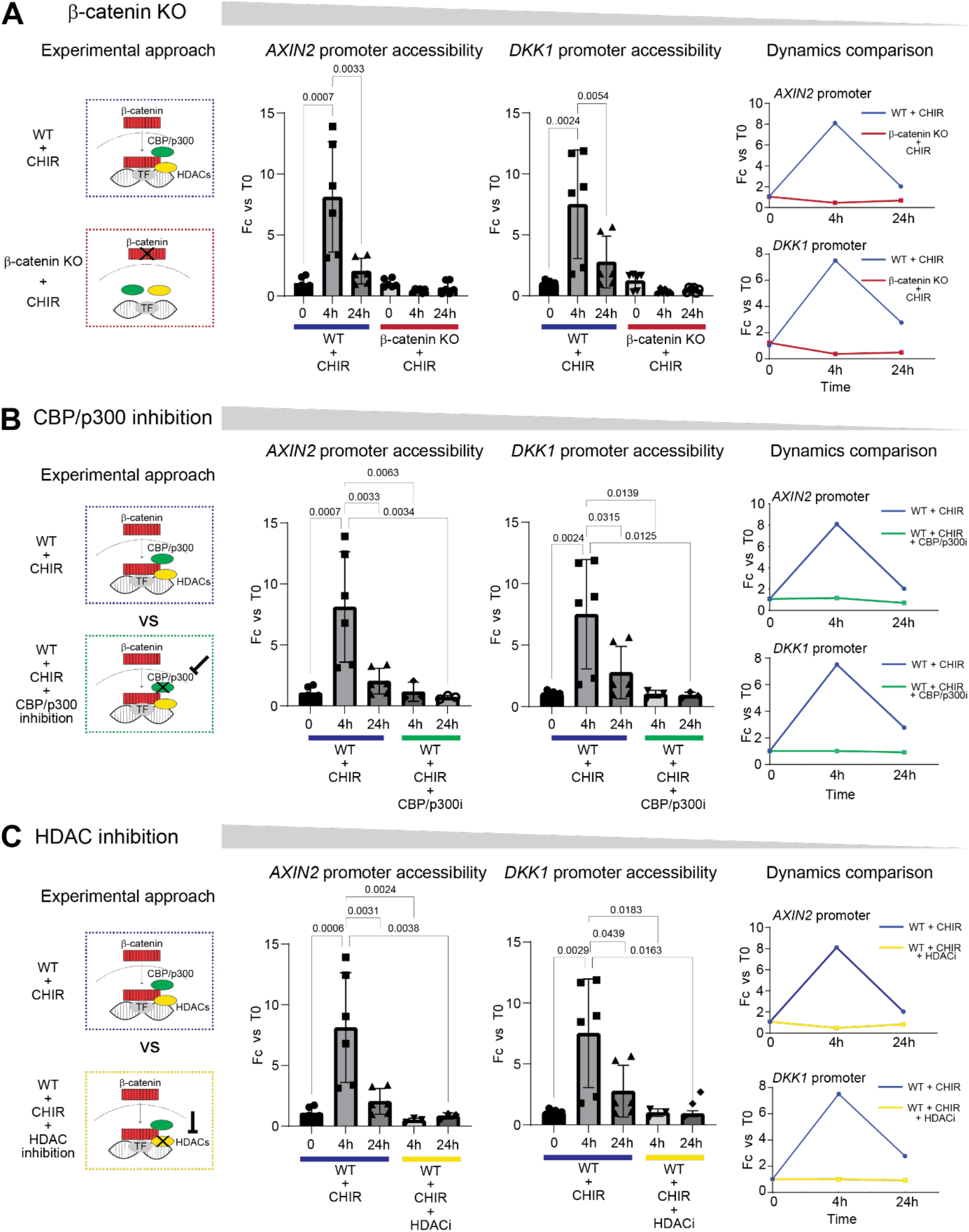
β-catenin drives chromatin remodeling at WREs via CBP/p300 and HDAC enzymatic activities. (A) Schematic representation of the β-catenin nuclear import, which follows CHIR administration (left). We treated parental wild-type and β-catenin KO HEK293T with CHIR (“WT + CHIR” and “β-catenin KO + CHIR”, respectively) and quantified chromatin accessibility at the promoters of the Wnt target genes *AXIN2* and *DKK1* by ATACqPCR. CHIR treatment alone recapitulated the elastic response described in Figure 5. β-catenin KO HEK293T cells failed to open chromatin at the *AXIN2* and *DKK1* promoters in response to CHIR stimulation (central and right plots). ATAC-qPCR values are expressed as fold-change against the respective time 0 value (WT, T0 for WT samples; KO, T0 for KO samples). (B) Schematic representation of the CBP/p300 and HDAC1 positioning within the β-catenin transcriptional complex according to the literature (left). Wild-type HEK293T cells were treated with CHIR and the CBP/p300 inhibitor A-485 (“WT + CHIR + CBP/p300 inhibition”) for 24 hours. Quantification of chromatin accessibility at the promoters of the Wnt target genes *AXIN2* and *DKK1* by qPCR (central and right panels). The concomitant treatment of HEK293T with CHIR and A-485 prevented chromatin opening at 4 hours and 24 hours. (C) Wild-type HEK293T cells were treated with CHIR and the HDAC inhibitor sodium butyrate (“WT + CHIR + HDAC inhibition”). Quantification of chromatin accessibility was measured as above. Sodium butyrate-mediated HDAC inhibition prevented CHIR-induced opening of the chromatin at 4 and 24 hours (central and right panels). Replicates: n = 6 for “WT + CHIR” samples, for each time point; n = 3 for all other samples. Samples were always compared one-vs-one using the One-way ANOVA test. The exact p-value is indicated on the plots; non-significant p-values (p > 0.05) are omitted. Abbreviations: Fc: fold change; HAT: histone acetyltransferase; HDAC: histone deacetylase; WT: wild type.

Nucleosomal histone acetylation is thought to be a pivotal mechanism in modulating chromatin accessibility (Klemm et al., 2019). We reasoned that, among the first β-catenin co-factors to be identified are the histone acetyl transferase (HAT) CBP/p300 and the histone deacetylase (HDAC1) chromatin-remodeling enzymes (Billin et al., 2000; Hecht et al., 2000). We tested their requirement for the β-catenin-induced chromatin opening, by exploiting the pharmacological inhibitory agents A-485 and sodium butyrate, which specifically and efficiently inhibit CBP/p300 (Lasko et al., 2017) and HDAC (Davie, 2003), respectively. The concomitant treatment with CHIR and either A-485 or sodium butyrate in parental HEK293T (namely, in the presence of functional β-catenin) fully abolished the CHIR-induced, β-catenin-dependent chromatin opening, as shown by the flattening of the response curves (Figure 6B, C). It is notable to observe that, while functional inhibition of CBP/p300 – HAT activity led to an outcome that is consistent with a previous report (Parker et al., 2008), the requirement of HDAC activity for the CHIR/β-catenin-mediated induction of chromatin opening was unanticipated.

Taken together, these analyses mechanistically show that the chromatin dynamics occurring upon CHIR stimulation require the presence of β-catenin, together with the chromatin remodeling activities of CBP/p300 and HDAC1 associated to it. This emphasizes two key new findings: i) the dynamic interplay between HAT and HDAC activities, rather than HAT alone, is at the basis of increased chromatin accessibility upon Wnt induction; ii) β-catenin, likely via its direct recruitment of CBP/p300 and HDAC1, possesses features that are typically associated with pioneer factors, and is responsible for the quantitative increase of chromatin accessibility observed upon CHIR treatment at target promoters.

## Discussion

The classic model of transcriptional activation of Wnt signaling is centered on the notion that stabilized β-catenin associates with the TCF/LEF family of transcription factors, on WREs, to induce the expression of Wnt target genes. However, this model does not provide a fulfilling explanation to three questions which, we believe, are key to understanding how cells respond to Wnt signaling: i) What is the temporal unfolding of the genome-wide binding behavior of β-catenin? ii) To what extent do different cell types present varying responses upon Wnt stimulation? iii) How is the chromatin modulated during this response on a global scale? To address these questions, here we employed a multi-level omics approach at select time points and resolved the response to Wnt/β-catenin activation in two paradigmatic human cell types. Our study includes the generation of a time-resolved roadmap of β-catenin physical chromatin occupancy over time, the comparison of the time-specific β-catenin binding events with the transcriptional responses, and how the chromatin is globally shaped by all these mechanisms with temporal resolution.

Only a few datasets have been previously produced with CUT&RUN-related technologies (i.e., mostly ChIP-seq) to identify the physical targets of β-catenin, due to the difficulty in detecting the behavior of this protein (Zambanini et al., 2022). By their own experimental design, these must be interpreted as snapshots of targets occurring at an arbitrarily chosen moment after stimulation. On the contrary, our setup reveals that the definition of Wnt target genes can be established only in the light of the temporal resolution. The first key observation apparent from our temporal CUT&RUN-LoV-U analysis is in fact that β-catenin “moves” to different targets along stimulation time, and its positioning occurs in proximity of genes whose action is consistent with the changing behavior of the cells considered. For example, β-catenin associated to pluripotency genes in the early moments of the response in hESCs, to then switch to lineage-specific ones at later time-points (Figure 2C-E). This shows that the regulation of targets genes does not occur in one momentous action that follows the signal but is diluted over time, likely to gradually foster cell behavior and lineage progression. Admittedly, the varying identity of targeted regions over time opens the possibility that our dataset is also limited, as it relies on the *a priori* choice of arbitrarily selected time-points. We posit that future experiments, with increased time-resolution or perhaps single cell approaches, will clarify this conundrum and bring new light into how individual cells shape their genomic response to signaling pathways.

Another pressing question that remains to be solved is what drives β-catenin repositioning over time. Time-resolved motif analysis of our datasets indicated that the answer might lie in the promiscuous cooperativity between β-catenin and other transcription factors (Figure 2E). In support to this, a recent report proposed that the stemness factor SOX2 determines β-catenin genomic localization in hESCs (Blassberg et al., 2022). This is consistent with the observed prominent association in our study between β-catenin peaks and motifs of SOX factors, enriched in particular at the early time points after Wnt activation in hESCs. We suspect that Blasberg and colleagues might have identified a piece of a more complex puzzle, whereby β-catenin localization changes over time, since after the interplay with SOX2, it engages with lineage-specific temporal partners. Our dataset supports this view, as the enrichment of SOX footprints is followed by the switch from SOX/OCT motifs with those for lineage-determining players, such as GATA4, required for mesoderm-derived cell fates (Molkentin et al., 1997) (Figure 2E). 20 years of research have been and are still exposing a growing body of literature identifying instances in which β-catenin cooperates with other transcription factors depending on the tissue (Söderholm and Cantù, 2020). Here, we propose that β-catenin promiscuity additionally occurs over time, and that its varying “affairs” might define the differentiation trajectories rather than being simply tissue-specific idiosyncrasies.

Striking was also the low overlap of physical β-catenin target genes between the two cell types used. HEK293T cells appear to modulate their response in a manner that is almost exclusively executed by the β-catenin-TCF/LEF complex, while no statistical enrichment for SOX and other pluripotency factors was observed. Of note, in HEK293T, the presence of the caudal-related homeodomain (CDX) motifs supports the identification by the Cadigan lab of a TCF/CDX module within a recently identified potent enhancer downstream of *AXIN2* (Ramakrishnan et al., 2021) (Figure 1D). We propose that the association with cell-specific transcription factors will explain in part the differential distribution of β-catenin in different models. Whether this fully depends on the distinctive association of β-catenin with tissue specific molecular partners (Blassberg et al., 2022; Zimmerli et al., 2020), by its recruitment to MED1-marked molecular condensates (Zamudio et al., 2019), or a combination of the two, remains to be established.

An open problem in the Wnt field of research is if there are reliable direct targets that become transcriptionally repressed by this pathway (Cadigan and Ramakrishnan, 2017). Few instances have been identified in the past, mostly by using *Drosophila melanogaster*, where the mechanisms were shown to be either interference with the binding of other transcription factors (Piepenburg et al., 2000), by alternative protein complexes with transcriptional repressors, such as Brinker (Theisen et al., 2007), or even due to the sequence specificity underlying regulatory modules of TCF/LEF on the DNA (Kim et al., 2017; Zhang et al., 2014). Our work generates a catalogue of the potential negatively regulated direct targets in these two cell types, by merging the direct binding position of β-catenin on regulatory regions over time (Figures 1-2) with the subsequent up- or down-regulation of target mRNAs (Figure 3). It came as a surprise that, upon pathway stimulation, the group sizes of repressed and activated genes associated with β-catenin peaks are comparable, possibly underscoring that the phenomenon of Wnt-mediated repression is more consistent than previously thought. Whether in human cells the mechanisms of repressions are the same as in the fruit fly remains to be established. Worth mentioning in this context is the recent functional association between the action of the transcription factor SP5, a upregulated target of Wnt signaling itself, and the repression of the Wnt-driven genetic programs in human pluripotent progenitor cells (Huggins et al., 2017). It is plausible that the observed downregulation constitutes a rapid, yet secondary response mediated by SP5 during the progression and termination of the Wnt-driven developmental program.

According to the Wnt-activation > SP5 repression model, the dual nature of positive or negative regulation of transcription could occur longitudinally, over time, rather than in distinct cells. This might be pertinent to the elastic chromatin response observed here (Figures 5-6). While β-catenin brings the chromatin remodelers CBP/p300 and HDAC1 to target loci, other factors, among which potentially SP5, could enter the scene in a timely fashion to promptly terminate this response. The recent identification of TBX3 as tissue-specific component of the Wnt/β-catenin transcriptional complex might also bring light to this mechanism (Zimmerli et al., 2020). In this study, TBX3 displayed an ambivalent mode of action combining enhanced repression or activation depending on the intensity of pathway stimulation. As *SP5, TBX3* is also a Wnt target gene mainly described as repressor (Renard et al., 2007; Willmer et al., 2017), and could constitute a transcriptional switch to modulate the response to Wnt in time. Consistent with this interpretation, *TBX3* is among the top upregulated direct targets specifically in HEK293T, where the elastic response is observed (Figure 3, Figure S2).

Finally, our work identifies that different genomes can respond in a dramatically divergent manner to the same extra-cellular stimulus. Overall, the chromatin accessibility profiles of HEK293T and hESCs are diverse before the stimulation, a conspicuous indicator of their distinct identities. One possible prediction could have been that Wnt/β-catenin, by activating homogeneous genetic programs – consisting of the upregulation of the same groups of genes – might have reduced the inter-cell type variability. Our datasets are suggestive of the opposite phenomenon: by acting on genomes with different chromatin characteristics, Wnt/β-catenin unequivocally increases cell type diversity by driving distinctive fates. hESCs, which *in vivo* activate the driver of mesoderm *TBXT*/*BRA* upon Wnt/β-catenin activation, do so also *in vitro*, to then gradually gain a mesodermal genetic program (Figures 3-4). HEK293T, on the other hand, more mildly fluctuate between neural and renal identities, consistent with previous reports (Lin et al., 2014). The relatively minor transcriptional changes associated with β-catenin peaks across time in HEK293T is indicative of these cells’ refractory tendency in changing cell identity. We speculate that the β-catenin-driven and CBP/p300-HDAC-mediated elastic chromatin response observed on a plethora of Wnt target genes in HEK293T might be causative of their inability of progressing toward other lineage identities. If further work will confirm this model, a clarifying picture might emerge, in which developmental Wnt/β-catenin signaling is assisted by tissue-specific transcription factors that operate progressive lineage choices. On the other hand, homeostatic Wnt signaling, which acts to preserve specific cell features, such as the stemness of *LGR5*^+^ cell in the intestinal epithelium (Sato et al., 2011) or the proliferative potential of cells close to the central vein in the liver (Jin et al., 2022), could generate a short-term, elastic response that drives the expression of groups of genes required to sustain that specific function, locally and in a tightly-regulated temporal manner.

## Acknowledgments

The authors are grateful to the laboratories of Prof. Francisca Lottersberger and Prof. Stefan Koch for continuous scientific input and reagents exchange. This work was supported by Grants to C.C. from Cancerfonden (CAN 2018/542 and 21 1572 Pj), and the Swedish Research Council, Vetenskapsrådet (2021-03075). C.C. is a Wallenberg Molecular Medicine (WCMM) fellow and receives generous financial support from the Knut and Alice Wallenberg Foundation. P.P is supported by a fellowship awarded by the WCMM at Linköping University. Computations and data handling were enabled by resources provided by the Swedish National Infrastructure for Computing (SNIC) at [SNIC CENTRE] partially funded by the Swedish Research Council through grant agreement no. 2018-05973.

## Author contributions

P.P. performed the experiments and prepared the figures. S.S. analyzed the ATAC- and RNA sequencing data and generated several figure components. A.N. and G.Z. performed and analyzed CUT&RUN-LoV-U. A.J.M. assisted with cell culture treatments and differentiation protocols. C.C supervised the research team and provided financial support for the study. P.P. and C.C. designed the projects, supervised the experimental execution, interpreted the data, and wrote the manuscript “with four hands”. All authors reviewed and commented on the final manuscript.

## Supplementary Figures

**Figure S1.**
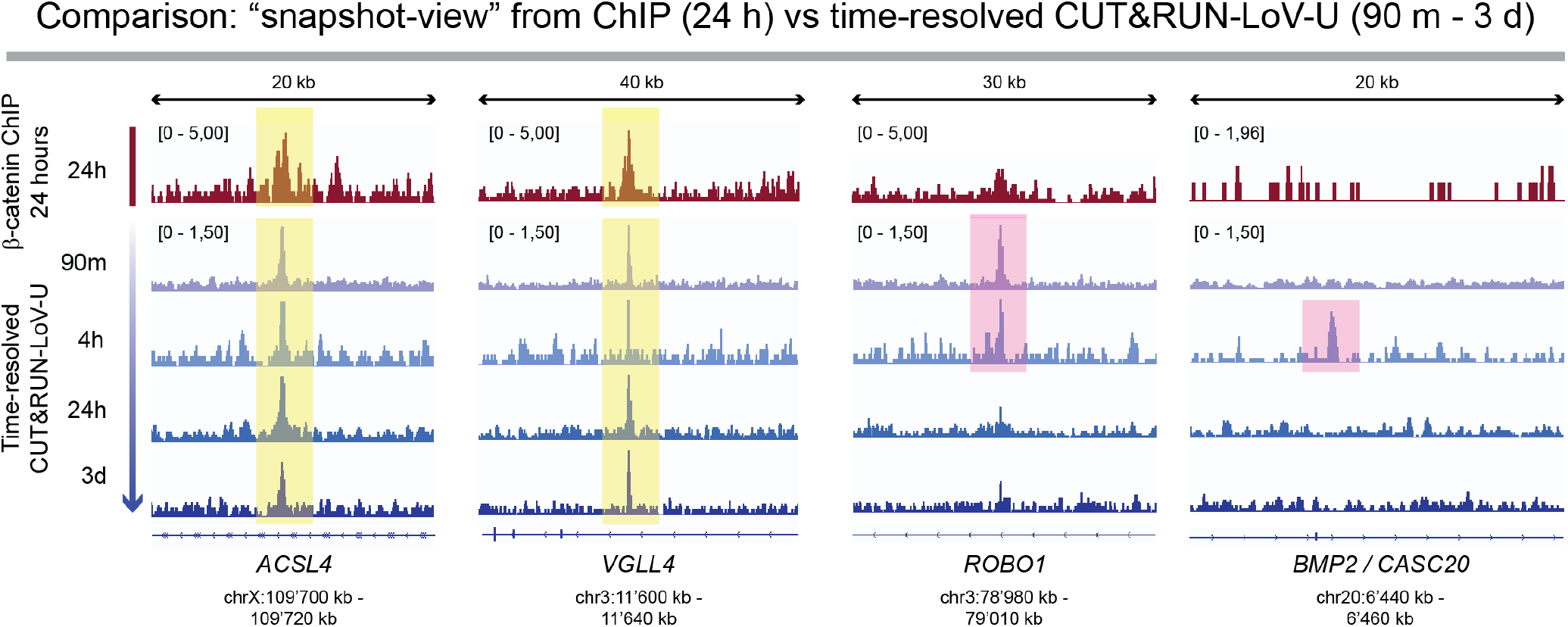
Comparison between single-time-point ChIP-seq and time-resolved CUT&RUN-LoV-U analysis of β-catenin genome-wide binding patterns. Time-resolved CUT&RUN-LoV-U performed at 90 minutes, 4 hours, 24 hours and, 3 days of CHIR stimulation (violet>blue tracks) robustly reproduces β-catenin binding patterns observed via Chromatin Immunoprecipitation-Sequencing (ChIP-seq, performed after 24 hours of CHIR stimulation, red track, dataset from Doumpas et al., 2019) at stable target loci (left panels, peaks highlighted by yellow boxes). However, single-time-point (“snapshot-view”) ChIP-seq failed to detect time-restricted β-catenin binding events (right panels, peaks highlighted by pink boxes).

**Figure S2.**
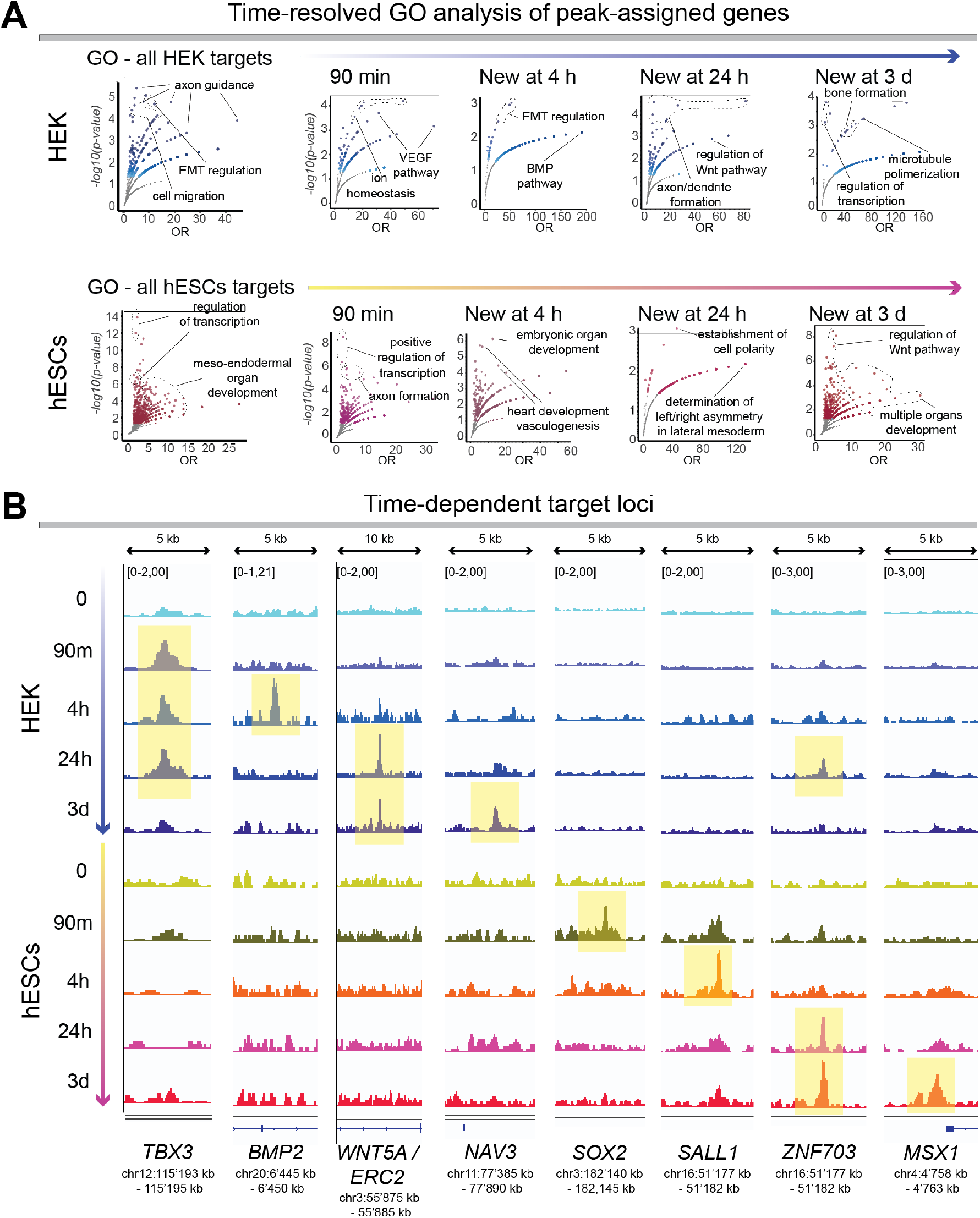
β-catenin target genes change along Wnt activation time. (A) Gene ontology (GO) analysis of all β-catenin physical targets (left dot-plots) and of β-catenin targets appearing at different time points following Wnt activation (dot-plots, from left to right). Selected GO terms are shown. First row: β-catenin targets in HEK293T; second row: β-catenin targets in hESCs. Genes associated with β-catenin peaks in HEK293T and hESCs are associated with different biological processes. Similarly, even within the same cells, time-specific β-catenin target genes mediate diverging processes over time. (B) Examples of time-specific β-catenin physical targets. *TBX3* is targeted specifically in HEK293T cells, starting 90 minutes after Wnt/β-catenin activation. *BMP2* is targeted solely in HEK293T at 4 hours. A region assigned to *WNT5A* is bound by β-catenin only from 24 hours of Wnt activation and onwards, while *NAV3* is targeted only after 3 days. Similar strong time-dependency was observed in hESCs. A putative *SOX2* regulatory region, located > 200 kb downstream of its 3’, is transiently but reproducibly bound by β-catenin at 90 minutes. In an analogous manner, a putative *SALL1* regulatory region is targeted at 4 hours only. The Wnt target gene *ZNF703* is bound by β-catenin in both HEK293T and hESCs at 24 hours; this β-catenin peak is maintained at 3 days in hESCs but lost in HEK293T. *MSX1* is targeted only in hESCs and only after 3 days of Wnt activation.

**Figure S3.**
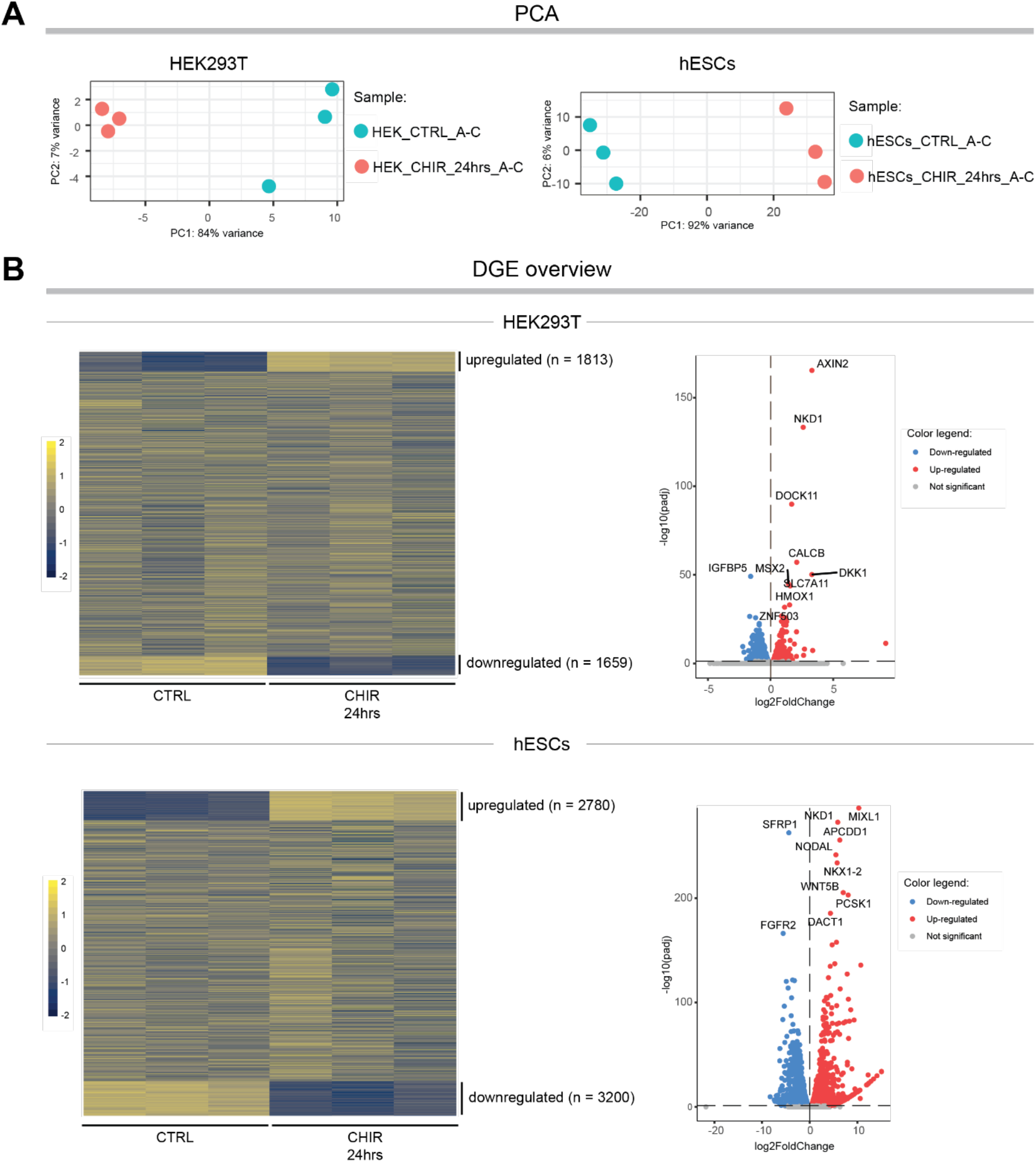
RNA sequencing overview. (A) Principal component analysis (PCA) correctly groups samples based on treatment. (B) Overview of differential gene expression induced by CHIR treatment in HEK293T (Doumpas et al., 2019) and hESCs (this work, see STAR Methods section). 1813 genes were upregulated, and 1659 genes were downregulated in HEK293T cells after 24 hours of CHIR treatment. The same treatment led to the upregulation of 2780 genes and the downregulation of 3200 genes in hESCs. Threshold for significance: padj < 0.05 (see STAR Methods section).

**Figure S4.**
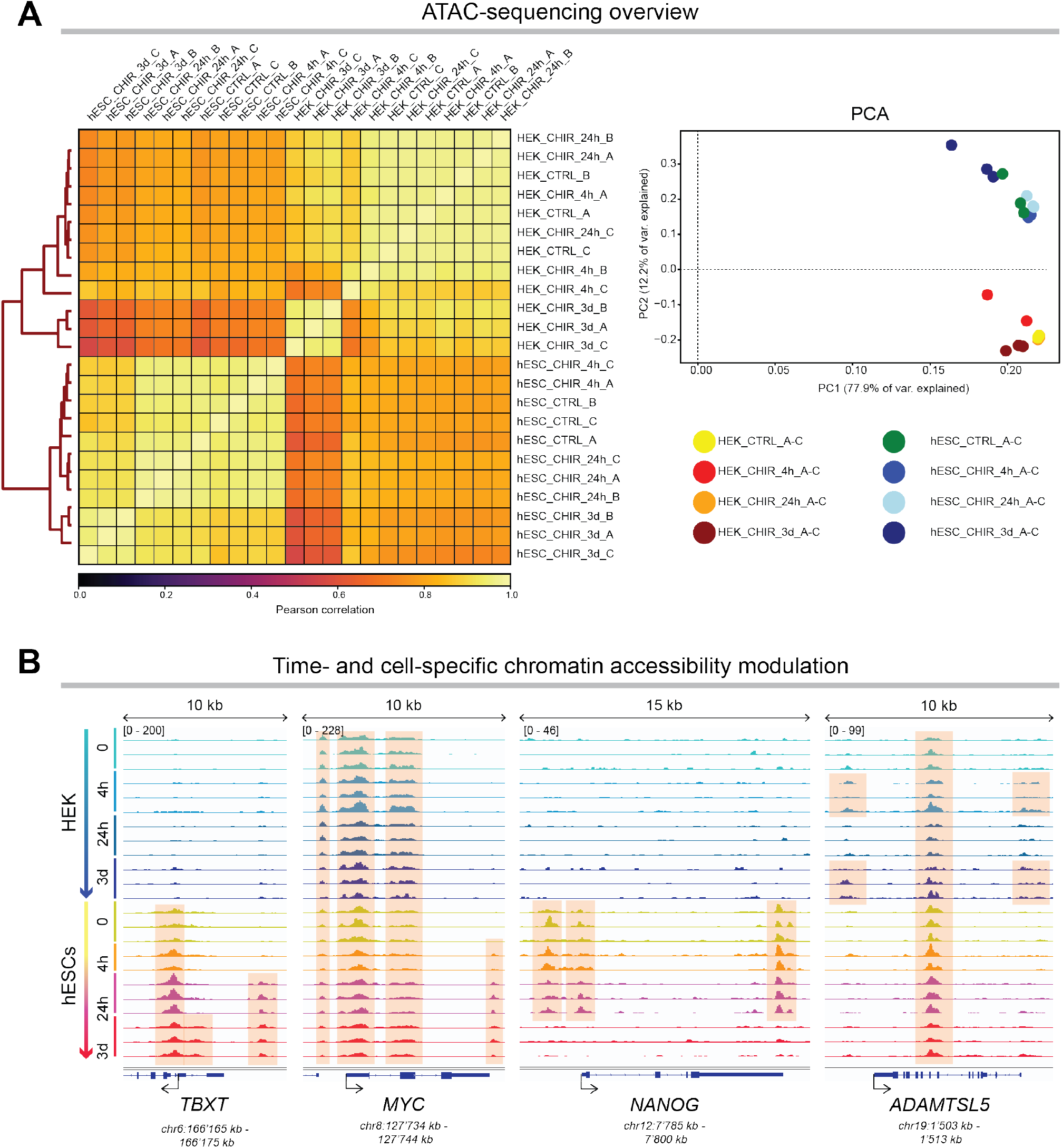
ATAC-sequencing overview. (A) Pearson correlation and Principal Component Analysis (PCA) of the time-resolved ATAC-sequencing dataset correctly group samples based on cell identity and treatment. (B) Examples of time- and cell-specific chromatin responses to Wnt/β-catenin signaling activation. hESCs show progressive opening of chromatin regions surrounding the mesodermal-differentiation-associated *TBXT* gene, as well as that of the key regulator of proliferation and global transcription *MYC*; in a complementary fashion, hESCs progressively close the chromatin regions that include and surround pluripotency factors, such as *NANOG*. HEK293T also shows time-specific alterations in chromatin accessibility, as exemplified by the *ADAMTSL5* locus.

**Figure S5.**
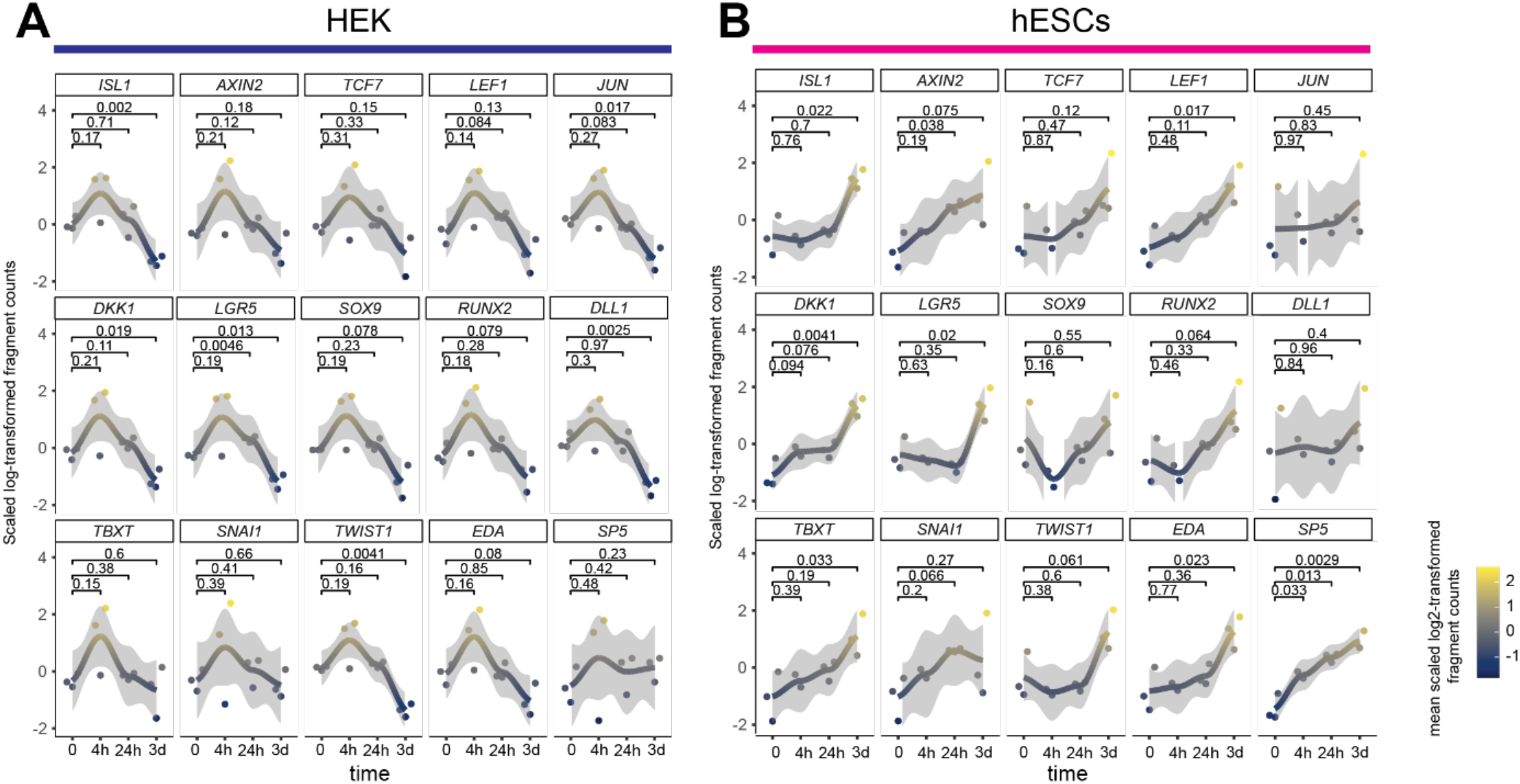
Elastic (HEK293T) and plastic (hESCs) chromatin responses at target loci. Line-plots representing differential chromatin accessibility profiles in correspondence with the promoter regions of all the individual targets analyzed. (A) Single chromatin accessibility profiles individually recapitulate almost invariably the elastic response in HEK293T. (B) The changing chromatin profiles in hESCs all display similar plastic trends as they progressively display increased accessibility over time.

## Methods

### Experimental models and subject details

Human embryonic stem cells (hESCs) were kindly provided by the group of Robert Passier (Den Hartogh et al., 2015). Human embryonic kidney (HEK) 293T cells belong to our laboratory (Doumpas et al., 2019).

### Data availability statement

All datasets described in this work are publicly available at ArrayExpress (https://www.ebi.ac.uk/arrayexpress/) with accession numbers E-MTAB-12077 (CUT&RUN-LoV-U), E-MTAB-12076 (ATAC-sequencing), and E-MTAB12075 (RNA-sequencing).

### Method details

#### Cell culture

HEK293T cells (HEK293Ts) were cultured in medium consisting of DMEM, high glucose (41966-029, Gibco – Thermo Fisher Scientific) supplemented with 10% Calf Bovine Serum (12133C, Sigma-Aldrich) and 10 U/ml Penicillin-Streptomycin (15276355, Gibco – Thermo Fisher Scientific). HEK293Ts were passaged by incubation in Trypsin EDTA 0.25% (25-200-056, Thermo Fisher Scientific). hESCs were cultured in Essential 8™ medium (A1517001, ThermoFisher Scientific) in plates and flasks coated with 10 μg/ml Vitronectin (A14700, ThermoFisher Scientific). Cells were passaged by incubation in 0.5 mM EDTA-PBS (stock: 0.5 M EDTA, AM9260G, Thermo Fisher Scientific).

#### Time resolved CUT & RUN for β-catenin

HEK293Ts were grown in culture medium supplemented with 10 nM LGK (S7143, Selleck Chemicals) for the 48 hours preceding the seeding. HEK293Ts were then seeded at a density of 10^4^ cells/cm^2^ in culture medium (see “Cell Culture” section) supplemented with 10 nM LGK. After 24 hours, cells were cultured in medium supplemented with 10 μM CHIR99021 (SML1046, Sigma Aldrich). Medium and treatment were renewed every 24 hours. hESCs were seeded at a density of 10^4^ cells/cm^2^ in Essential 8™ medium, and after 24 hours, they were cultured in Essential 8™ medium supplemented with 10 μM CHIR99021. Cells were then detached (as described in the “Cell Culture” section) after 90 minutes, 4 hours, 24 hours, and 72 hours of CHIR treatment. All experiments were performed in triplicate (n = 3 for each cell type at each time-point). Cells were filtered through a 40 μm cell strainer (KKE3.1, Carl Roth) and processed for CUT&RUN (Zambanini et al., 2022). 500,000 cells/sample were harvested using either Trypsin EDTA 0.25% (HEK293Ts) or EDTA 0.5 mM in PBS (hESCs) and washed twice in DPBS (Cat. #14190094, Thermo Fisher Scientific). Cells were washed three times in Nuclear Extraction (NE) buffer (HEPES-KOH pH-8.2 [20 mM], KCl [10 mM], Spermidine [0.5 mM], IGEPAL [0.05%], Glycerol [20%], Roche Complete Protease Inhibitor EDTA-Free), then resuspended in 40 μl NE per sample and bound to 20 μl Magnetic ConA Agarose beads equilibrated in binding buffer. After incubation, nuclei and beads were resuspended for 5 minutes in EDTA wash buffer (wash buffer with EDTA [0.2 mM]). Samples were divided in 200 μl PCR tubes and antibody incubation was performed in 200 μl wash buffer with 2 μl of anti-β-catenin antibody (Cat. #ABIN2855042, antibodies-online) overnight (ON) at 4 °C on a rotator. After ON incubation, samples were washed five times in wash buffer and resuspended in 200 μl of pAG-MN buffer (wash buffer supplemented with pAG-MN [0.6 μg/ml]) for 30 minutes at 4 °C on a rotator. Samples were washed five times, followed by digestion for 30 minutes in wet ice in wash buffer with 2 mM CaCl2. After 30 minutes, the digestion buffer was removed and the reaction was stopped with 50 μl of 1X Urea STOP buffer (NaCl [100 mM], EDTA [2 mM], EGTA [2 mM], IGEPAL [0.5%], Urea [8.8 M]) and the samples were incubated for 1 hour at 4°C. Beads were collected on the magnetic rack, while the liquid was transferred to a new PCR tube, where it was cleaned up twice using Mag-Bind TotalPure NGS beads (Cat. #M1327, Omega BioTek) at 2X, and then resuspended in 20 μl Tris-HCl pH 7.5.

##### Library preparation and sequencing

Library preparation was performed using the KAPA Hyper Prep Kit for Illumina platforms (Cat. #KK8504, KAPA Biosystems) according to the manufacturer’s guidelines with the following modifications. End repair and A-tailing was performed with 0.4 size reactions with 20 µl of purified DNA. The thermocycler conditions were set to 12 °C for 15 min, 37 °C for 15 min and 58 °C for 25 min to prevent thermal degradation of the shortest fragments. Adapter ligation was done with 0.4 size reactions. KAPA Dual Indexed adapters were used at 0.15 µM. A post-ligation clean-up was performed with Mag-Bind TotalPure NGS beads at 1.2X. Resuspension was done in 10 mM Tris-HCl pH 8.0. Library amplification was performed with 0.5 size reactions. The cycling conditions were set as follows: 45 sec initial denaturation at 98 °C, 15 sec denaturation at 98 °C, 10 sec annealing/elongation at 60 °C, 1 min final extension at 72 °C, hold at 4 °C, with 13 cycles. After amplification, a post-amplification cleanup was performed with 1.2X beads. Libraries were then run on an E-Gel EX 2% agarose gel (Cat. # G402022, Invitrogen) for 10 min using the E-Gel Power Snap Electrophoresis System (Invitrogen). Bands of interest between 150 and 500 bp were cut out and purified using the GeneJET Gel Extraction Kit (Cat. #K0691, Thermo Scientific) according to manufacturer’s instructions. Libraries were quantified with the Qubit (Thermo Scientific) using their high sensitivity DNA kit (Cat #Q32854, Thermo Scientific), pooled and sequenced 36 bp pair-end on the NextSeq 550 (Illumina) using the Illumina NextSeq 500/550 High Output Kit v2.5 (75 cycles) (Cat. #20024906, Illumina). The CUT&RUN datasets (raw and processed files) have been deposited at ArrayExpress (https://www.ebi.ac.uk/arrayexpress/) under accession number E-MTAB-12077.

##### Data Analysis

Read quality was assessed using fastqc (Brandine & Smith, 2022, version 0.11.9). Trimming was performed using bbmap bbduk (Bushnell *et al*, 2017, version 38.18) removing adapters, artifacts, poly AT, G and C repeats. One replicate of HEK293T Chir 4 hours was removed from downstream analysis due to low read count and poor quality. Reads were aligned to the hg38 genome with bowtie (Langmead *et al*, 2009, version 1.0.0) using options -v 0 -m 1 -X 500. Samtools (Li et al., 2009), version 1.11) view, fixmate, markdup and sort were used to create bam files, mark and remove duplicates, and sort bam files. Individual track bedgraphs were created using bedtools (Quinlan & Hall, 2010, version 2.23.0) genomecov on pair-end mode. BAM files of biological replicates were merged using samtools and mitochondrial reads were removed. Normalized signal per million reads tracks for visualization were created by using the --SPRM function of macs2 (Zhang *et al*, 2008, version 2.2.6) for each replicate pool with the options -f BAMPE --keep-dup all --SPMR and –bdg. Merged BAM files were randomly shuffled and split into two pseudoreplicates. Peaks were called using macs2 for each pseudoreplicate against the corresponding negative control using the options -f BAMPE –keep-dup-all and -p 1e-2. Narrowpeak output files were sorted by p value, and input into IDR (doi:10.1214/11-AOAS466, version 2.0.4.2) with the options –rank p.value and –idr-threshold 0.2 to determine high confidence peaks for each condition. Each negative control file was peak called separately with macs2 with the options -f BAMPE –keep-dup-all and -q 5e-2. Final peak sets for each condition were generated by using bedtools subtract with the option -A to remove blacklisted regions (Amemiya et al., 2019) and regions overlapping with peaks called in the corresponding negative control. Venn diagrams and overlap peak sets were created using Intervene (Khan & Mathelier, 2017, version 0.6.4). Motif analysis was done using Homer (Heinz *et al*, 2010, version 4.11) findMotifsGenome to find motifs in the hg38 genome using -size given. Peak set gene annotation was done using GREAT (McLean *et al*, 2010, version 4.0.4) with default parameters. Signal intensity plots were created using ngsplot (Shen *et al*, 2014, version 2.63) with options -N 2 -GO hc -SC global.

#### Time-resolved ATAC-sequencing

##### Cells treatment, library preparation and sequencing

HEK293Ts and hESCs were seeded as described in the “Time resolved CUT & RUN for β-catenin” section. Both cell types were treated with 10 μM CHIR99021 for 4 hours, 24 hours and 3 days (n = 3 samples for each condition). At the end of the culture period, 10^4^ cells were processed for ATAC-sequencing. ATAC-sequencing was performed according to previously published protocols (Buenrostro et al., 2016). Briefly, 10^4^ cells were collected, lysed in ice cold lysis buffer (Tris-HCl pH 7.4 [10 mM], NaCl [10 mM], MgCl2 [3 mM], IGEPAL CA-630 [0.1%]), and pelleted. Pelleted nuclei were incubated in 50 μl transposition reaction mix (20034210, Illumina) at 37^0^C for 30 minutes. Transposed DNA fragments were amplified for 13 cycles in the presence of Custom Nextera PCR primers (Buenrostro et al., 2013) using the NEBNext High-Fidelity 2x PCR Master Mix (Cat. #M0541, New England Biolabs). Libraries were purified using the High Pure PCR Production Purification Kit (11732676001, Roche / Sigma-Aldrich). Libraries were validated on an Agilent 2100 and quantified using quantitative PCR (Q-PCR). Libraries were then sequenced on Illumina NovaSeq 6000 S4 flow-cell with PE150 according to results from library quality control and expected data volume. The ATAC-seq datasets (raw and processed files) have been deposited at ArrayExpress under accession number E-MTAB-12076.

##### Quality control, pre-processing, and alignment

The sequencing quality of the obtained fastq data files was assessed using the quality control tool FastQC (version 0.11.5) with default settings. Furthermore, the tool FastQ Screen ((Wingett and Andrews, 2018), version 0.13.0) was used to ensure the correct species origin of the starting material, and all quality control results were summarized using multiQC ((Ewels et al., 2016), version 1.7). Sequencing reads were trimmed to remove adapter contamination and to filter out reads with a sequencing quality score below 30, using the bbduk tool from the BBTools suite (version 38.58) in paired-end mode. Trimming parameters were “ktrim=r k=23 mink=11 hdist=1 tpe tbo” for adapter removal and “qtrim=l trimq=30” for quality filtering. Trimmed reads were mapped to the human reference genome GRCh38 using the alignment tool Bowtie2 ((Langmead and Salzberg, 2012), version 2.3.4.1) in paired-end mode and with –very-sensitive-local setting. Pre-built bowtie index files excluding ALT contigs were downloaded from the bowtie2 page at SourceForge (http://bowtie-bio.sourceforge.net/bowtie2/in-dex.shtml). Resulting sequence alignment map (SAM) files were converted to binary alignment map (BAM) files, sorted by coordinates and indexed using samtools ((Danecek et al., 2021), version 1.9).

##### Data overview and genome track visualization

To get a first overview of the differences and similarities between the sequenced samples the data was processed using tools from the Deeptools suite ((Ramírez et al., 2016), version 3.3.0). Scaling factors were computed with the multiBamSummary tool, which were then used for sequencing depth normalization during read coverage computation using the bamCoverage tool. Duplicated reads, reads mapped to the mitochondrial chromosome, and reads overlapping blacklisted regions (as defined by the ENCODE project, data downloaded from https://github.com/Boyle-Lab/Blacklist/blob/master/lists/hg38-blacklist.v2.bed.gz) were removed. Read coverage data was saved in bigwig format and visualized as genomic tracks using the Integrative Genome Viewer (IGV, version 2.4.17, (Robinson et al., 2011)). Next, to investigate the overall similarity of the different bigwig files, the average scores in every genomic region across the genome were computed using multiBigwigSummary, with default bin size of 10kb. The results were used to calculate sample correlation (Pearson correlation method with outliers removed) and preform principal component analysis (PCA).

##### Peak calling

Genomic regions enriched for mapped reads over background read coverage was identified using the peak calling tool Genrich (version 0.6), taking sample replicates into consideration and removing duplicated reads and reads in blacklisted regions. A Benjamini-Hochberg false discovery rate (FDR)-adjusted p-value (q) < 0.05 and minimum area under the curve (AUC) of 200 were used as thresholds for statistical significance. Called peaks were further analyzed using the R programming language (version 4.1.0) and the developmental environment software Rstudio (version 1.1.463). Peaks were annotated with genomic information using the R package ChIPseeker ((Yu et al., 2015), version 1.28.3). Peak co-localization across samples was assessed through peak overlap analysis using the R package ChIPpeakAnno ((Zhu et al., 2010), version 3.26.4), where peaks were considered overlapping if at least one base pair was shared. Genes associated with differential peaks were then analyzed with the Gene Ontology (Biological Processes) online tool (Ashburner et al., 2000; Carbon et al., 2021; Mi et al., 2019).

##### Differential transcription factor accessibility analysis

Called peaks of chromatin accessibility were analyzed for differential transcription factor (TF) accessibility using the Differential ATAC-seq tool kit (DAStk, version 1.0.0, (Tripodi et al., 2018)) with default settings. Pre-scanned TF motif files from the TF model database HOCOMOCO (Kulakovskiy et al., 2018) were downloaded from https://github.com/Dowell-Lab/DAStk (Kulakovskiy et al., 2018). In short, each peak file was processed to calculate motif displacement (MD) scores which were then compared between samples to produce differential MD scores and corresponding p-values for each TF motif. A threshold of p < 1e-5 was used for determining statistical significance.

##### Plastic/Elastic chromatin accessibility response for selected regions

Peak regions at the promoter of a set of known WNT target genes (WTGs) were selected through visual inspection of the ATAC-seq signal tracks in IGV. Mapped reads were assigned to these selected regions using the read summarization tool FeatureCounts ((Liao et al., 2014), version 2.0.1). Fragments, rather than the reads themselves, were counted. Only fragments with a minimum length of 20bp and a maximum length of 120bp were included while multimapped reads were excluded. These count data were visualized in heatmaps using the pheatmap R package, applying scaling to the data for proper comparison between the time points. Bar plots were generated with GraphPad Prism (version 9.3.0).

#### RNA sequencing

RNA sequencing was performed on RNA isolated from 5*10^5^ hESCs cells obtained from the same wells used for the time-resolved ATAC-sequencing analysis. Total RNA was extracted with the RNeasy Mini Kit (Cat. #74106, Qiagen). 500 ng of total RNA were processed with the Illumina Stranded mRNA Prep kit (Cat. #20040898, #20040896, #20040899, #20026121, Illumina) for library preparation. Messenger RNA (mRNA) was purified from total RNA using poly-T oligo-attached magnetic beads. After fragmentation, the first strand cDNA was synthesized using random hexamer primers followed by the second strand cDNA synthesis. cDNA was then subjected to end repair, A-tailing, adapter ligation, size selection, amplification, and purification. Libraries were then sequenced on Illumina NovaSeq 6000 S4 flow-cell with PE150 according to results from library quality control and expected data volume. These RNA-seq datasets (raw and processed files) have been deposited at ArrayExpress under accession number E-MTAB-12075. For HEK293T cells, we used the dataset published by Doumpas et al., 2019.

##### Data analysis

Quality control of the raw data was performed as described in the “Time-resolved ATAC-sequencing” section. Read quality and adapter contamination trimming was done with the bbduk tool from the BBTools suite (version 38.58) in paired-end mode, with parameters “ktrim=l k=23 mink=11 hdist=1 qtrim=rl trimq=25 minlength=10 trimpolyg=10 tpe tbo”. The Spliced Transcript Alignment to a Reference (STAR) software (version 2.7.3a) (Dobin et al., 2013) was used to map pre-processed sequencing reads to the GRCh38.p13 human reference genome. STAR was run in paired-end mode with --quantMode GeneCounts settings. The reference genome (primary assembly) in FASTA format and associated primary assembly gene annotation in GTF format were downloaded from GENCODE, release 40 (https://www.gencodegenes.org/human/release_40). Differential gene expression analysis was performed using DESeq2 (version 1.32.0) (Love et al., 2014) with default settings. A Benjamini-Hochberg false discovery rate (FDR)-adjusted p-value below 0.05 was used as threshold for statistical significance.

#### Analysis of β-catenin KO HEK293T cells, and CBP/p300 and HDAC inhibition

To investigate the role of β-catenin in CHIR-induced chromatin responses, we used β-catenin KO HEK293T cells we generated previously (Doumpas et al., 2019). β-catenin KO HEK293T cells were grown in culture medium supplemented with 10 nM LGK (S7143, Selleck Chemicals) for the 48 hours preceding the seeding. HEK293Ts were then seeded at a density of 10^4^ cells/cm^2^ in culture medium (see “Cell Culture” section) supplemented with 10 nM LGK. After 24 hours, cells were cultured in medium supplemented with 10 μM CHIR99021 (SML1046, Sigma Aldrich). Medium and treatment were renewed every 24 hours. To investigate the role of CBP/p300 and HDACs in Wnt/β-catenin dependent chromatin responses, wild-type HEK293T cells were grown in culture medium supplemented with 10 nM LGK (S7143, Selleck Chemicals) for the 48 hours preceding the seeding. HEK293Ts were then seeded at a density of 10^4^ cells/cm^2^ in culture medium (see “Cell Culture” section) supplemented with 10 nM LGK. After 24 hours, cells were cultured in medium supplemented with 10 μM CHIR99021 (SML1046, Sigma Aldrich) and 5 μM A-485 (CBP/p300 inhibitor; (Lasko et al., 2017)) or 1 mM Sodium Butyrate (broad HDAC inhibitor) for 4 hours and 24 hours. At the end of the culture period, 5*10^4^ cells were processed for ATAC-qPCR (cells were treated, and their DNA tagmented, as described in the “Time-resolved ATAC-sequencing” section).

##### ATAC-qPCR

The tool ATACPrimerTool (Yost et al., 2018) was used to predict optimal regions for ATAC-qPCR primers based on the ATAC-seq data. Sorted bam files were used as input and the analysis was performed with default settings. ATAC-qPCR quantification of chromatin accessibility in the different conditions was normalized on the values obtained from WNT-OFF conditions. All analyses were performed on at least n = 3 independent replicates (the number of replicates for each experiment is indicated in the figure legends), from 5*10^4^ cells / replicate, processed in parallel and amplified for the same number of cycles during ATAC-DNA-library preparation. The following primers were used: *AXIN2*promoter: Fw - CCA GGA CCT TAT CAA AGC GC, Rv - GGA TCA CTG GCT CCG CGA; *DKK1*promoter: Fw - TGC TAT AAC GCT CGC TGG TA, Rv - GGA TGG GAT TTC AAA GCG CT. Samples were always compared one-vs-one using the One-way ANOVA test (Graph Pad Prism 9.3.0). The exact p-value is indicated on the plots.

## References

Agbleke, A.A., Amitai, A., Buenrostro, J.D., Chakrabarti, A., Chu, L., Hansen, A.S., Koenig, K.M., Labade, A.S., Liu, S., Nozaki, T., et al. (2020). Advances in Chromatin and Chromosome Research: Perspectives from Multiple Fields. Mol. Cell 79, 881–901.

Aksoy, I., Giudice, V., Delahaye, E., Wianny, F., Aubry, M., Mure, M., Chen, J., Jauch, R., Bogu, G.K., Nolden, T., et al. (2014). Klf4 and Klf5 differentially inhibit mesoderm and endoderm differentiation in embryonic stem cells. Nat. Commun. 2014 51 5, 1–15.

Amemiya, H.M., Kundaje, A., and Boyle, A.P. (2019). The ENCODE Blacklist: Identification of Problematic Regions of the Genome. Sci. Rep. 9, 1–5.

Ashburner, M., Ball, C.A., Blake, J.A., Botstein, D., Butler, H., Cherry, J.M., Davis, A.P., Dolinski, K., Dwight, S.S., Eppig, J.T., et al. (2000). Gene Ontology: tool for the unification of biology. Nat. Genet. 25, 25.

Baker, J.C., Beddington, R.S.P., and Harland, R.M. (1999). Wnt signaling in Xenopus embryos inhibits Bmp4 expression and activates neural development. Genes Dev. 13, 3149–3159.

Billin, A.N., Thirlwell, H., and Ayer, D.E. (2000). β-Catenin–Histone Deacetylase Interactions Regulate the Transition of LEF1 from a Transcriptional Repressor to an Activator. Mol. Cell. Biol. 20, 6882–6890.

Blassberg, R., Patel, H., Watson, T., Gouti, M., Metzis, V., Delás, M.J., and Briscoe, J. (2022). Sox2 levels regulate the chromatin occupancy of WNT mediators in epiblast progenitors responsible for vertebrate body formation. Nat. Cell Biol. 24.

Brandine, G.D.S., and Smith, A.D. (2022). Falco : high-speed FastQC emulation for quality control of sequencing data [version 2 ; peer review : 2 approved].

Buenrostro, J., Wu, B., Chang, H., and Greenleaf, W. (2016). ATAC-seq: A Method for Assaying Chromatin Accessibility Genome-Wide. Curr. Protoc. Mol. Biol. 190, 21.29.1-21.29.9.

Buenrostro, J.D., Giresi, P.G., Zaba, L.C., Chang, H.Y., and Greenleaf, W.J. (2013). Transposition of native chromatin for fast and sensitive epigenomic profiling of open chromatin, DNA-binding proteins and nucleosome position. Nat. Methods 10, 1213–1218.

Bushnell, B., Rood, J., and Singer, E. (2017). BBMerge –Accurate paired shotgun read merging via overlap. 1–15.

Cadigan, K.M., and Ramakrishnan, A.B. (2017). Wnt target genes and where to find them. F1000Research 6, 746.

Cantù, C., Felker, A., Zimmerli, D., Prummel, K.D., Cabello, E.M., Chiavacci, E., Méndez-Acevedo, K.M., Kirchgeorg, L., Burger, S., Ripoll, J., et al. (2018). Mutations in Bcl9 and Pygo genes cause congenital heart defects by tissue-specific perturbation of Wnt/β-catenin signaling. Genes Dev. 32, 1443–1458.

Carbon, S., Douglass, E., Good, B.M., Unni, D.R., Harris, N.L., Mungall, C.J., Basu, S., Chisholm, R.L., Dodson, R.J., Hartline, E., et al. (2021). The Gene Ontology resource: Enriching a GOld mine. Nucleic Acids Res. 49, D325–D334.

Chang, J.L., Chang, M. V., Barolo, S., and Cadigan, K.M. (2008a). Regulation of the feedback antagonist naked cuticle by Wingless signaling. Dev. Biol. 321, 446–454.

Chang, M. V., Chang, J.L., Gangopadhyay, A., Shearer, A., and Cadigan, K.M. (2008b). Activation of Wingless Targets Requires Bipartite Recognition of DNA by TCF. Curr. Biol. 18, 1877–1881.

Cole, M.F., Johnstone, S.E., Newman, J.J., Kagey, M.H., and Young, R.A. (2008). Tcf3 is an integral component of the core regulatory circuitry of embryonic stem cells. Genes Dev. 22, 746–755.

Danecek, P., Bonfield, J.K., Liddle, J., Marshall, J., Ohan, V., Pollard, M.O., Whitwham, A., Keane, T., McCarthy, S.A., Davies, R.M., et al. (2021). Twelve years of SAMtools and BCFtools. Gigascience 10.

Davidson, K.C., Adams, A.M., Goodson, J.M., McDonald, C.E., Potter, J.C., Berndt, J.D., Biechele, T.L., Taylor, R.J., and Moon, R.T. (2012). Wnt/β-catenin signaling promotes differentiation, not self-renewal, of human embryonic stem cells and is repressed by Oct4. Proc. Natl. Acad. Sci. U. S. A. 109, 4485–4490.

Davie, J.R. (2003). Inhibition of Histone Deacetylase Activity by Butyrate. J. Nutr. 133, 2485–2493.

Dias, A., Lozovska, A., Wymeersch, F.J., Nóvoa, A., Binagui-Casas, A., Sobral, D., Martins, G.G., Wilson, V., and Mallo, M. (2020). A Tgfbr1/Snai1-dependent developmental module at the core of vertebrate axial elongation. Elife 9, 1–28.

Doumpas, N., Lampart, F., Robinson, M.D., Lentini, A., Nestor, C.E., Cantù, C., and Basler, K. (2019). TCF / LEF dependent and independent transcriptional regulation of Wnt/β-catenin target genes. EMBO J. 38, e98873.

Doumpas, N., Söderholm, S., Narula, S., Moreira, S., Doble, B.W., Cantù, C., and Basler, K. (2021). TCF/LEF regulation of the topologically associated domain ADI promotes mESCs to exit the pluripotent ground state. Cell Rep. 36.

Efroni, S., Duttagupta, R., Cheng, J., Dehghani, H., Hoeppner, D.J., Dash, C., Bazett-Jones, D.P., Le Grice, S., McKay, R.D.G., Buetow, K.H., et al. (2008). Global Transcription in Pluripotent Embryonic Stem Cells. Cell Stem Cell 2, 437–447.

Ewels, P., Magnusson, M., Lundin, S., and Käller, M. (2016). MultiQC: summarize analysis results for multiple tools and samples in a single report. Bioinformatics 32, 3047–3048.

Fafilek, B., Krausova, M., Vojtechova, M., Pospichalova, V., Tumova, L., Sloncova, E., Huranova, M., Stancikova, J., Hlavata, A., Svec, J., et al. (2013). Troy, a tumor necrosis factor receptor family member, interacts with lgr5 to inhibit wnt signaling in intestinal stem cells. Gastroenterology 144, 381–391.

Gujral, T.S., and Macbeath, G. (2010). A System-Wide Investigation of the Dynamics of Wnt Signaling Reveals Novel Phases of Transcriptional Regulation. PLoS One 5, e10024.

Den Hartogh, S.C., Schreurs, C., Monshouwer-Kloots, J.J., Davis, R.P., Elliott, D.A., Mummery, C.L., and Passier, R. (2015). Dual Reporter MESP1mCherry/w-NKX2-5eGFP/w hESCs Enable Studying Early Human Cardiac Differentiation. Stem Cells 33, 56–67.

Hecht, A., Vleminckx, K., Stemmler, M.P., Roy, F. van, and Kemler, R. (2000). The p300/CBP acetyltransferases function as transcriptional coactivators of β-catenin in vertebrates. EMBO J. 19, 1839–1850.

Heinz, S., Benner, C., Spann, N., Bertolino, E., Lin, Y.C., Laslo, P., Cheng, J.X., Murre, C., Singh, H., and Glass, C.K. (2010). Simple Combinations of Lineage-Determining Transcription Factors Prime cis -Regulatory Elements Required for Macrophage and B Cell Identities. Mol. Cell 38, 576–589.

Hinrichs, A.S., Karolchik, D., Baertsch, R., Barber, G.P., Bejerano, G., Clawson, H., Diekhans, M., Furey, T.S., Harte, R.A., Hsu, F., et al. (2006). The UCSC Genome Browser Database : update 2006. 34, 590–598.

Huggins, I.J., Bos, T., Gaylord, O., Jessen, C., Lonquich, B., Puranen, A., Richter, J., Rossdam, C., Brafman, D., Gaasterland, T., et al. (2017). The WNT target SP5 negatively regulates WNT transcriptional programs in human pluripotent stem cells. Nat. Commun. 2017 81 8, 1–14.

Irion, U., and Nüsslein-Volhard, C. (2022). Developmental Genetics of Model Organisms. Proc. Natl. Acad. Sci. U. S. A. 119, 117–117.

Jho, E., Zhang, T., Domon, C., Joo, C.-K., Freund, J.-N., and Costantini, F. (2002). Wnt/β-Catenin/Tcf Signaling Induces the Transcription of Axin2, a Negative Regulator of the Signaling Pathway. Mol. Cell. Biol. 22, 1172–1183.

Jiao, S., Li, C., Hao, Q., Miao, H., Zhang, L., Li, L., and Zhou, Z. (2017). VGLL4 targets a TCF4-TEAD4 complex to coregulate Wnt and Hippo signalling in colorectal cancer. Nat. Commun. 8.

Jin, Y., Anbarchian, T., Wu, P., Sarkar, A., Fish, M., Chuan, W., and Nusse, R. (2022). Wnt signaling regulates hepatocyte cell division by a transcriptional repressor cascade. Proc. Natl. Acad. Sci. U. S. A. 1–9.

van Kappel, E.C., and Maurice, M.M. (2017). Molecular regulation and pharmacological targeting of the β-catenin destruction complex. Br. J. Pharmacol. 174, 4575–4588.

Kelly, K.F., Ng, D.Y., Jayakumaran, G., Wood, G.A., Koide, H., and Doble, B.W. (2011). β-catenin enhances Oct-4 activity and reinforces pluripotency through a TCF-independent mechanism. Cell Stem Cell 8, 214–227.

Khan, A., and Mathelier, A. (2017). Intervene: A tool for intersection and visualization of multiple gene or genomic region sets. BMC Bioinformatics 18, 1–8.

Kim, K., Cho, J., Hilzinger, T.S., Nunns, H., Liu, A., Ryba, B.E., and Goentoro, L. (2017). Two-Element Transcriptional Regulation in the Canonical Wnt Pathway. Curr. Biol. 27, 2357-2364.e5.

Klemm, S.L., Shipony, Z., and Greenleaf, W.J. (2019). Chromatin accessibility and the regulatory epigenome. Nat. Rev. Genet. 20, 207–220.

Kulakovskiy, I. V., Vorontsov, I.E., Yevshin, I.S., Sharipov, R.N., Fedorova, A.D., Rumynskiy, E.I., Medvedeva, Y.A., Magana-Mora, A., Bajic, V.B., Papatsenko, D.A., et al. (2018). HOCOMOCO: towards a complete collection of transcription factor binding models for human and mouse via large-scale ChIP-Seq analysis. Nucleic Acids Res. 46, D252–D259.

Kuleshov, M. V, Jones, M.R., Rouillard, A.D., Fernandez, N.F., Duan, Q., Wang, Z., Koplev, S., Jenkins, S.L., Jagodnik, K.M., Lachmann, A., et al. (2016). Enrichr : a comprehensive gene set enrichment analysis web server 2016 update. 44, 90–97.

de la Roche, M., and Bienz, M. (2007). Wingless-independent association of Pygopus with dTCF target genes. Curr. Biol. 17, 556–561.

Langmead, B., and Salzberg, S.L. (2012). Fast gapped-read alignment with Bowtie 2. Nat. Methods 9, 357–359.

Langmead, B., Trapnell, C., Pop, M., and Salzberg, S.L. (2009). Ultrafast and memory-efficient alignment of short DNA sequences to the human genome. 10.

Lasko, L.M., Jakob, C.G., Edalji, R.P., Qiu, W., Montgomery, D., Digiammarino, E.L., Hansen, T.M., Risi, R.M., Frey, R., Manaves, V., et al. (2017). Discovery of a selective catalytic p300/CBP inhibitor that targets lineage-specific tumours. Nature 550, 128–132.

Li, H., Handsaker, B., Wysoker, A., Fennell, T., Ruan, J., Homer, N., Marth, G., Abecasis, G., and Durbin, R. (2009). The Sequence Alignment/Map format and SAMtools. Bioinformatics 25, 2078–2079.

Li, V.S.W., Ng, S.S., Boersema, P.J., Low, T.Y., Karthaus, W.R., Gerlach, J.P., Mohammed, S., Heck, A.J.R., Maurice, M.M., Mahmoudi, T., et al. (2012). Wnt Signaling through Inhibition of β-Catenin Degradation in an Intact Axin1 Complex. Cell 149, 1245–1256.

Liao, Y., Smyth, G.K., and Shi, W. (2014). featureCounts: an efficient general purpose program for assigning sequence reads to genomic features. Bioinformatics 30, 923–930.

Lin, Y.C., Boone, M., Meuris, L., Lemmens, I., Van Roy, N., Soete, A., Reumers, J., Moisse, M., Plaisance, S., Drmanac, R., et al. (2014). Genome dynamics of the human embryonic kidney 293 lineage in response to cell biology manipulations. Nat. Commun. 5.

MacDonald, B.T., and He, X. (2012). Frizzled and LRP5/6 Receptors for Wnt/?-Catenin Signaling. Cold Spring Harb. Perspect. Biol. 4, a007880–a007880.

Mähler, M., Berar, M., Feinstein, R., Gallagher, A., Illgen-Wilcke, B., Pritchett-Corning, K., and Raspa, M. (2014). FELASA recommendations for the health monitoring of mouse, rat, hamster, guinea pig and rabbit colonies in breeding and experimental units. Lab. Anim. 48, 178–192.

Marks, H., Kalkan, T., Menafra, R., Denissov, S., Jones, K., Hofemeister, H., Nichols, J., Kranz, A., Francis Stewart, A., Smith, A., et al. (2012). The transcriptional and epigenomic foundations of ground state pluripotency. Cell 149, 590–604.

Martin, B.L., and Kimelman, D. (2010). Brachyury establishes the embryonic mesodermal progenitor niche. Genes Dev. 24, 2778–2783.

McLean, C.Y., Bristor, D., Hiller, M., Clarke, S.L., Schaar, B.T., Lowe, C.B., Wenger, A.M., and Bejerano, G. (2010). GREAT improves functional interpretation of cis-regulatory regions. Nat. Biotechnol. 28, 495–501.

Meers, M.P., Bryson, T.D., Henikoff, J.G., and Henikoff, S. (2019a). Improved CUT&RUN chromatin profiling tools. Elife 8, 1–16.

Meers, M.P., Tenenbaum, D., and Henikoff, S. (2019b). Peak calling by Sparse Enrichment Analysis for CUT&RUN chromatin profiling. Epigenetics and Chromatin 12, 1–11.

Mi, H., Muruganujan, A., Ebert, D., Huang, X., and Thomas, P.D. (2019). PANTHER version 14: more genomes, a new PANTHER GO-slim and improvements in enrichment analysis tools. Nucleic Acids Res. 47, D419–D426.

Molkentin, J.D., Lin, Q., Duncan, S.A., and Olson, E.N. (1997). Requirement of the transcription factor GATA4 for heart tube formation and ventral morphogenesis. Genes Dev. 11, 1061–1072.

Moon, R.T., and Kimelman, D. (1998). From cortical rotation to organizer gene expression: Toward a molecular explanation of axis specification in Xenopus. BioEssays 20, 536–546.

Moreira, S., Seo, C., Polena, E., Mahendram, S., Mercier, E., Blais, A., and Doble, B.W. (2018). TCF7L1 and TCF7 differentially regulate specific mouse ES cell genes in response to GSK-3 inhibition. BioRxiv 1–14.

Mosimann, C., Hausmann, G., and Basler, K. (2009). Beta-catenin hits chromatin: regulation of Wnt target gene activation. Nat. Rev. Mol. Cell Biol. 10, 276–286.

Orkin, S.H., and Hochedlinger, K. (2011). Chromatin connections to pluripotency and cellular reprogramming. Cell 145, 835–850.

Parker, D.S., Ni, Y.Y., Chang, J.L., Li, J., and Cadigan, K.M. (2008). Wingless signaling induces widespread chromatin remodeling of target loci. Mol. Cell. Biol. 28, 1815–1828.

Piepenburg, O., Vorbrüggen, G., and Jäckle, H. (2000). Drosophila Segment Borders Result from Unilateral Repression of Hedgehog Activity by Wingless Signaling. Mol. Cell 6, 203–209.

Quinlan, A.R., and Hall, I.M. (2010). BEDTools: A flexible suite of utilities for comparing genomic features. Bioinformatics 26, 841–842.

Ramakrishnan, A.B., Chen, L., Burby, P.E., and Cadigan, K.M. (2021). Wnt target enhancer regulation by a CDX/TCF transcription factor collective and a novel DNA motif. Nucleic Acids Res. 49, 8625–8641.

Ramírez, F., Ryan, D.P., Grüning, B., Bhardwaj, V., Kilpert, F., Richter, A.S., Heyne, S., Dündar, F., and Manke, T. (2016). deepTools2: a next generation web server for deep-sequencing data analysis. Nucleic Acids Res. 44, W160–W165.

Reinhold, M.I., Kapadia, R.M., Liao, Z., and Naski, M.C. (2006). The Wnt-inducible Transcription Factor Twist1 Inhibits Chondrogenesis. J. Biol. Chem. 281, 1381–1388.

Renard, C.A., Labalette, C., Armengol, C., Cougot, D., Wei, Y., Cairo, S., Pineau, P., Neuveut, C., De Reyniès, A., Dejean, A., et al. (2007). Tbx3 is a downstream target of the Wnt/β-catenin pathway and a critical mediator of β-catenin survival functions in liver cancer. Cancer Res. 67, 901–910.

Rim, E.Y., Clevers, H., and Nusse, R. (2022). The Wnt Pathway: From Signaling Mechanisms to Synthetic Modulators. Annu. Rev. Biochem. 91, 1–28.

Robinson, J.T., Thorvaldsdóttir, H., Winckler, W., Guttman, M., Lander, E.S., Getz, G., and Mesirov, J.P. (2011). Integrative genomics viewer. Nat. Biotechnol. 29, 24–26.

Roux, M., Laforest, B., Capecchi, M., Bertrand, N., and Zaffran, S. (2015). Hoxb1 regulates proliferation and differentiation of second heart field progenitors in pharyngeal mesoderm and genetically interacts with Hoxa1 during cardiac outflow tract development. Dev. Biol. 406, 247–258.

Sato, T., van Es, J.H., Snippert, H.J., Stange, D.E., Vries, R.G., van den Born, M., Barker, N., Shroyer, N.F., van de Wetering, M., and Clevers, H. (2011). Paneth cells constitute the niche for Lgr5 stem cells in intestinal crypts. Nature 469, 415–418.

Schuijers, J., Mokry, M., Hatzis, P., Cuppen, E., and Clevers, H. (2014). Wnt-induced transcriptional activation is exclusively mediated by TCF/LEF. EMBO J. 33, 146–156.

Shaw, G., Morse, S., Ararat, M., and Graham, F.L. (2002). Preferential transformation of human neuronal cells by human adenoviruses and the origin of HEK 293 cells. FASEB J. 16, 869–871.

Shen, L., Shao, N., Liu, X., and Nestler, E. (2014). ngs. plot : Quick mining and visualization of next-generation sequencing data by integrating genomic databases. 1–14.

Skene, P.J., Henikoff, J.G., and Henikoff, S. (2018). Targeted in situ genome-wide profiling with high efficiency for low cell numbers. Nat. Protoc. 13, 1006–1019.

Söderholm, S., and Cantù, C. (2020). The WNT/β-catenin dependent transcription: A tissue-specific business. WIREs Syst. Biol. Med. 1–41.

Tajonar, A., Maehr, R., Hu, G., Sneddon, J.B., Rivera-Feliciano, J., Cohen, D.E., Elledge, S.J., and Melton, D.A. (2013). VGLL4 is a novel regulator of survival in human embryonic stem cells. Stem Cells 31, 2833–2841.

Theisen, H., Syed, A., Nguyen, B.T., Lukacsovich, T., Purcell, J., Srivastava, G.P., Iron, D., Gaudenz, K., Nie, Q., Wan, F.Y.M., et al. (2007). Wingless Directly Represses DPP Morphogen Expression via an Armadillo/TCF/Brinker Complex. PLoS One 2, e142.

van Tienen, L.M., Mieszczanek, J., Fiedler, M., Rutherford, T.J., Bienz, M., Labhart, T., Desplan, C., Hursh, D., Jones, T., Bejsovec, A., et al. (2017). Constitutive scaffolding of multiple Wnt enhanceosome components by Legless/BCL9. Elife 6, 477–488.

Tripodi, I.J., Allen, M.A., and Dowell, R.D. (2018). Detecting Differential Transcription Factor Activity from ATAC-Seq Data. Molecules 23.

Weidinger, G., Thorpe, C.J., Wuennenberg-Stapleton, K., Ngai, J., and Moon, R.T. (2005). The Sp1-Related Transcription Factors sp5 and sp5-like Act Downstream of Wnt/β-Catenin Signaling in Mesoderm and Neuroectoderm Patterning. Curr. Biol. 15, 489–500.

Van de Wetering, M., Cavallo, R., Dooijes, D., Van Beest, M., Van Es, J., Loureiro, J., Ypma, A., Hursh, D., Jones, T., Bejsovec, A., et al. (1997). Armadillo coactivates transcription driven by the product of the Drosophila segment polarity gene dTCF. Cell 88, 789–799.

Wiese, K.E., Nusse, R., and van Amerongen, R. (2018). Wnt signalling: Conquering complexity. Dev. 145, 1–9.

Willmer, T., Cooper, A., Peres, J., Omar, R., and Prince, S. (2017). The T-Box transcription factor 3 in development and cancer. Biosci. Trends 11, 254–266.

Wingett, S.W., and Andrews, S. (2018). FastQ Screen: A tool for multi-genome mapping and quality control. F1000Research 7, 1338.

Wray, J., Kalkan, T., Gomez-Lopez, S., Eckardt, D., Cook, A., Kemler, R., and Smith, A. (2011). Inhibition of glycogen synthase kinase-3 alleviates Tcf3 repression of the pluripotency network and increases embryonic stem cell resistance to differentiation. Nat. Cell Biol. 13, 838–845.

Ying, Q.L., Wray, J., Nichols, J., Batlle-Morera, L., Doble, B., Woodgett, J., Cohen, P., and Smith, A. (2008). The ground state of embryonic stem cell self-renewal. Nature 453, 519–523.

Yost, K.E., Carter, A.C., Xu, J., Litzenburger, U., and Chang, H.Y. (2018). ATAC Primer Tool for targeted analysis of accessible chromatin. Nat. Methods 15, 304–305.

Yu, G., Wang, L.G., and He, Q.Y. (2015). ChIPseeker: an R/Bioconductor package for ChIP peak annotation, comparison and visualization. Bioinformatics 31, 2382–2383.

Zabidi, M.A., Arnold, C.D., Schernhuber, K., Pagani, M., Rath, M., Frank, O., and Stark, A. (2015). Enhancer-core-promoter specificity separates developmental and housekeeping gene regulation. Nature 518, 556–559.

Zambanini, G., Nordin, A., Jonasson, M., Pagella, P., and Cantù, C. (2022). A New CUT & RUN Low Volume-Urea (LoV-U) protocol uncovers Wnt / beta-catenin tissue-specific genomic targets. BioRxiv 2022, 1–12.

Zamudio, A. V., Dall’Agnese, A., Henninger, J.E., Manteiga, J.C., Afeyan, L.K., Hannett, N.M., Coffey, E.L., Li, C.H., Oksuz, O., Sabari, B.R., et al. (2019). Mediator Condensates Localize Signaling Factors to Key Cell Identity Genes. Mol. Cell 76, 753-766.e6.

Zhang, C.U., Blauwkamp, T.A., Burby, P.E., and Cadigan, K.M. (2014). Wnt-Mediated Repression via Bipartite DNA Recognition by TCF in the Drosophila Hematopoietic System. PLoS Genet. 10.

Zhang, Y., Liu, T., Meyer, C.A., Eeckhoute, J., Johnson, D.S., Bernstein, B.E., Nussbaum, C., Myers, R.M., Brown, M., Li, W., et al. (2008). Model-based analysis of ChIP-Seq (MACS). Genome Biol. 9.

Zhu, L.J., Gazin, C., Lawson, N.D., Pagès, H., Lin, S.M., Lapointe, D.S., and Green, M.R. (2010). ChIPpeakAnno: a Bioconductor package to annotate ChIP-seq and ChIP-chip data. BMC Bioinformatics 11.

Zimmerli, D., Borrelli, C., Jauregi-Miguel, A., Söderholm, S., Brütsch, S., Doumpas, N., Reichmuth, J., Murphy-Seiler, F., Aguet, M., Basler, K., et al. (2020). TBX3 acts as tissue-specific component of the Wnt/β-catenin enhanceosome. Elife 1–17.

